# Scalable ARG-free Detection of Denisovan-mediated Superarchaic Introgression Reveals Heterogeneous Patterns across Populations

**DOI:** 10.64898/2026.06.25.734355

**Authors:** Noel McAllister, Sebastian Zöllner, Xinjun Zhang

**Author notes:** These authors contributed equally to this work (Sebastian Zöllner, Xinjun Zhang).

## Abstract

Ghost introgression from unsampled hominin lineages has emerged as an increasingly important component of human evolutionary history. Recent studies suggest that deeply divergent hominin lineages may have contributed ancestry either directly to modern humans or indirectly through Denisovan introgression, while inference remains difficult due to few reference genomes, weak signal, and uncertainty in reconstructing deep genealogies. Here we show analytically and through simulations that Denisovan-mediated superarchaic introgression produces predictable shifts in local coalescent depth that can be approximated by scalable summary statistics, particularly pairwise sequence divergence, suggesting that substantial information regarding deeply divergent ancestry is preserved in sequence variations without explicit reconstruction of genealogies. Leveraging this insight, we develop DEEP (**D**eep ancestry **E**stimation through **E**fficient **P**roxies), an ARG-free neural-network framework for identifying candidate regions of superarchaic ancestry. DEEP retains detectable power at low false positive rates across a broad range of demographic parameter space, remains scalable and recovers signals from small sample sizes. Applying DEEP to Oceanians, Tibetans, and Han Chinese, we identify approximately 0.4-0.6% of genomic windows with evidence of superarchaic ancestry. Candidate regions show both substantial overlap and notable heterogeneity across populations, with repeated enrichment near the HLA locus across all populations, suggesting immune-related regions recurrently retain deeply divergent ancestry.

## 1 Introduction

We are living in an exciting era for studying archaic introgression in human evolution. High-coverage genomes from Neanderthals and Denisovans, along with increasingly large modern human genomic datasets, have revealed that interbreeding with divergent hominin groups was widespread and recurrent throughout human evolutionary history [1–7]. Beyond the known archaic lineages represented by currently available ancient genomes, there is growing evidence that additional, deeply divergent “ghost” populations contributed to the ancestry of modern humans, either directly through ghost introgression or indirectly through Neanderthal or Denisovan lineages through the so-called superarchaic introgression [8–13]. Additionally, fossil records particularly in Asia were morphologically diverse and overlapped geographically and temporally with modern humans and Denisovans, implying that hominin interaction during the Middle to Late Pleistocene was potentially more complex than previously thought [14–18].

Detecting ghost or superarchaic introgression remains methodologically challenging. Unlike Neanderthal or Denisovans, ghost populations lack reference genomes and therefore cannot be identified by direct donor-recipient comparison [19–22]. Furthermore, introgressed segments from Neanderthals and Denisovans are relatively rare and highly fragmented due to recombination, negative selection, and drift acting over thousands of generations [6, 23–25]. If ghost ancestry was inherited indirectly through introgressing into Denisovans, their tracts are expected to be even lower in frequency and smaller in length. What further complicates the inference is that signals appearing to be of ghost ancestry can alternatively arise from alternative demographic scenarios, including ancestral population structure. Finally, for populations of interest such as the Oceanians who harbor the highest amount of Denisovan ancestry [7, 26–28], are incompletely understood in terms of demographic history, which makes robust inference difficult.

Recent methodological advances have begun to identify genomic signatures consistent with ghost or superarchaic introgression [11–13]. For example, Durvasula and Sankararaman provided foundational evidence for ghost introgression in Africans using reference-free local ancestry inference and site frequency statistics [9]. Collier et al. used two-locus summaries from individual unphased genomes and inferred recurrent hominin gene flow, including deeply divergent ancestry entering Denisovans [29]. More recently, Zhang et al. reconstructed genome-wide Ancestral Recombination Graphs (ARGs) and reported evidence of superarchaic ancestry in Oceanians [30]. At the same time, alternative models such as deep population structure have been proposed to explain some of the ghost signals [31–35]. For example, Ragsdale et al. showed that a weakly structured stem model can explain some apparent ghost signature in Africa [33], and Loya et al. found only limited support for superarchaic introgression using ARG-based analyses [36]. These findings showed both promises and uncertainty surrounding ghost introgression, including its magnitude, genomic distribution, donor origin, and robustness to demographic confounders. Nevertheless, evidence across multiple methodological frameworks converges on the possibility that deeply divergent ancestry contributed to Denisovan evolutionary history.

In principle, ghost introgression can be identified through unusually deep coalescence in local genealogies and elevated distribution of Time to Most Recent Common Ancestors (TMRCAs). Existing ARG-based approaches infer this signal by explicitly reconstructing local genealogies across the genome. However, inference of ARG remains computationally intensive, difficult to scale, and inconsistent in accuracy and output topologies [13, 37–41]. As a result, there is currently no scalable framework for identifying where ghost introgression resides in the genome and how it varies across populations. More fundamentally, reconstructing full ARGs may be unnecessary. The signal of Denisovan-mediated superarchaic introgression is expected to reside in only a small subset of local genealogical relationships, whereas most inferred branches are unrelated to the introgression event of interest. Consequently, accurate detection may require only a fraction of the information contained within a full ARG. Moreover, because ghost ancestry is expected to be rare and weak, uncertainty distributed across many reconstructed genealogical relationships could potentially obscure the true ghost introgression signal. Many of the expected effects of ghost introgression, including the unusually deep human-Denisovan divergence, may instead be preserved in observable sequence variations that approximate local genealogical relationship. This raises a broader question: how much information regarding ghost ancestry can be recovered without reconstructing the genealogies themselves?

In this work, rather than reconstructing complete genealogical histories, we ask whether the key signal of ghost ancestry, the unusually deep local coalescence, can be sufficiently captured by simpler and more scalable approximations. We show analytically and through simulations that Denisovan-mediated superarchaic introgression produces detectable shifts toward unusually deep coalescence and that much of this signal is preserved in summary statistics, particularly pairwise sequence divergence, without explicit ARG reconstruction. Leveraging this insight, we develop DEEP (**D**eep ancestry **E**stimation through **E**fficient **P**roxies), an ARG-free neural-network framework trained on population-specific demographic simulations to identify candidate regions of superarchaic ancestry in modern human genomes. DEEP is computationally scalable and in contrast to many approaches that benefit from large population-scale datasets, DEEP recovers biologically meaningful signals using only a modest number of sampled individuals per population, making it particularly useful for populations where large genomic datasets remain unavailable.

Applying DEEP to Oceanians, Tibetans and Han Chinese, we identify a sparse genomic landscape of superarchaic ancestry characterized by both substantial overlap and notable heterogeneity across populations, suggesting a complex combination of shared ancestry, population-specific evolutionary processes, and potentially heterogenous interactions between Denisovan subgroups and superarchaic species.

## 2 Results

### 2.1 Superarchaic introgression produces predictable shifts in local coalescent depth

Our inference strategy rests on a simple theoretical expectation: if a deeply diverged ghost lineage contributed ancestry to Denisovans, and Denisovans later contributed ancestry to modern humans, then some present-day human haplotypes should carry genomic segments whose ancestry follows the path Ghost →Denisovan →human. The key genealogical consequence of this history is an increase in local human–Denisovan coalescent depth, because the relevant lineage traces back toward the older ghost– Denisovan split rather than only to the human–Denisovan split [42–45]. The demographic model used for this analytical calculation is shown in Fig. 1a. The assumed data are local genotype windows containing 14 phased modern-human haplotypes. We also evaluated the same calculation after adding a Denisovan reference, represented as two Denisovan pseudo-haplotypes, to ask how much the archaic reference changes the expected genealogical contrast.

**Fig. 1:**
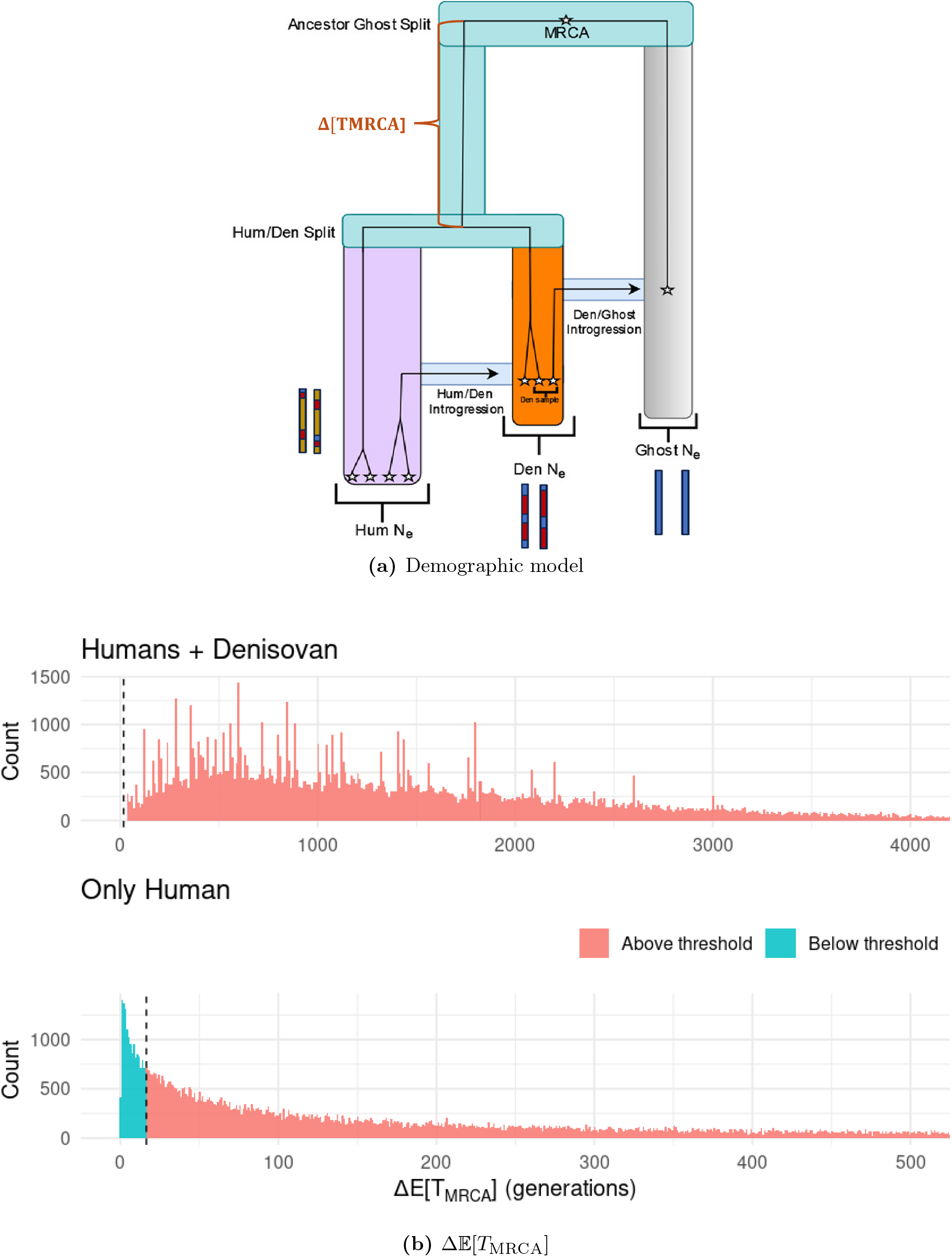
Analytical basis for detecting Ghost*→*Denisovan*→*human ancestry. **a**, Demographic model over which the analytical parameters are varied. A deeply diverged ghost lineage contributes ancestry to Denisovans, and Denisovans subsequently contribute ancestry to modern humans. The assumed observed data are local genomic windows containing 14 phased modern-human haplotypes; in the reference-inclusive calculation, two Denisovan pseudo-haplotypes are added. **b**, Distribution of the expected increase in coalescent depth under ghost ancestry relative to the no-ghost baseline. The top panel includes the modern-human haplotypes and two Denisovan pseudohaplotypes, whereas the bottom panel uses the modern-human haplotypes alone. The dashed vertical line gives the one-sided detectability threshold for 200 genomic windows. Values to the right of the threshold are parameter combinations whose expected coalescent-depth shift exceeds this heuristic cutoff.

Rather than evaluating a single parameterization of this model, we calculated the expectation over a grid of demographic and introgression parameters for the model in Fig. 1a. The grid varied the Denisovan →human ancestry fraction, the ghost → Denisovan ancestry fraction, the human–Denisovan and ghost–Denisovan split times, the two introgression pulse times, and the human and Denisovan effective population sizes. In broad terms, the grid spanned Denisovan → human ancestry fractions from 5% to 30%, ghost →Denisovan ancestry fractions from 1% to 5%, human–Denisovan split times from 300 to 900 ka, ghost–Denisovan split times from 1.0 to 1.8 Ma, pulse times from 40 to 700 ka, and effective population sizes from 1000 to 10,000. The full parameter grid and coalescent calculation are given in Methods.

For each parameter combination, we calculated the probability that sampled ancestry enters a mixed Denisovan-side state: at least one Denisovan-side lineage traces through the ghost lineage, while at least one Denisovan-side lineage remains outside the ghost contribution. This condition corresponds to 1 *≤ M*_G_ *< M*_D_, where *M*_D_ is the number of Denisovan-side lineages at the older pulse and *M*_G_ is the number of those lineages tracing through the ghost pulse. The strict inequality retains a contrast between ghost-derived and non-ghost Denisovan-side ancestry, rather than counting cases in which all Denisovan-side lineages have the same ghost ancestry.

Across the modern-human-only analytical grid, this mixed-state probability ranged from 1.5 × 10^−5^ to 0.106, with a median of 0.0058 and an interquartile range of 0.0017–0.0158. We then converted this probability into an expected change in coalescent depth, Δ*E*[*T*_MRCA_] = *E* [*T*_with ghost_] −*E* [*T*_no ghost_]. To summarize whether a parameter combination produced a shift large enough to be detectable in this analytical setting, we used a one-sided normal approximation. Specifically, we compared the expected shift to the standard error of an average over 200 windows under the no-ghost baseline (see Methods). We therefore use “detectable” to mean that the expected shift is larger than this one-sided normal-approximation cutoff. This is a heuristic scale comparison for the analytical grid, not a formal empirical test applied directly to observed genomic windows.

For the modern-human-only calculation, Δ*E* [*T*_MRCA_] ranged from 0.06 to 6388.0 generations. The median shift was 155.7 generations, with an interquartile range of 42.6–465.7 generations. Under a 25-year generation time, the median shift corresponds to approximately 3893 years.

Adding the Denisovan reference increased the expected shift in coalescent depth across the same parameter grid. With two Denisovan pseudo-haplotypes included, the median mixed-state probability increased to 0.0417, and the median expected shift increased to 1173.1 generations. In this reference-inclusive calculation, all parameter combinations exceeded the 200 window detectability threshold (Fig. 1b).

### 2.2 Detectability depends on demographic and introgression parameters

We next asked which parts of the parameter grid produced expected coalescent-depth shifts large enough to meet the detectability criterion described above and in Methods. These heatmaps use the modern-human-only calculation where we observed variability in detectability. They summarize the expected signal available from 14 phased modern-human haplotypes before adding a Denisovan reference. Across the full analytical grid, 100,410 of 115,200 parameter combinations, or 87.2%, exceeded the one-sided normal-approximation cutoff for 200 genomic windows.

Detectability increased with both introgression fractions (Fig. 2a). Across cells defined by the Denisovan → human and ghost → Denisovan ancestry fractions, the detectable proportion ranged from 35.7% to 99.7%. The lowest value occurred when the Denisovan → human ancestry fraction was 0.05 and the ghost→Denisovan ancestry fraction was 0.01-when both assumed their lowest values in the parameter grid. The highest value occurred when these fractions were 0.30 and 0.05 respectively, the maximum values of each in the parameter grid. The median expected shift across these cells increased from 11.3 generations in the weakest introgression cell to 973.9 generations in the strongest.

**Fig. 2:**
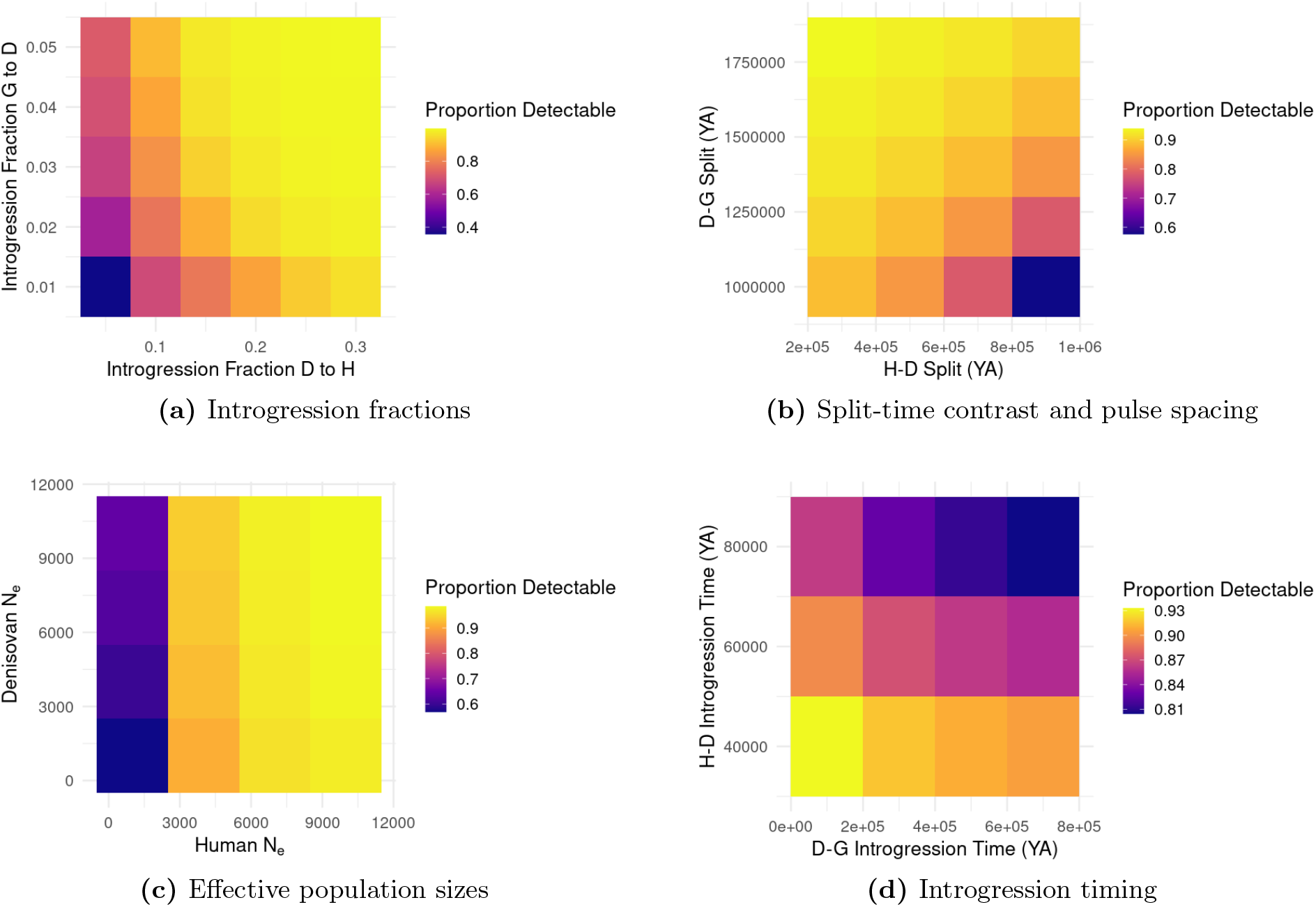
Detectability of Ghost*→*Denisovan*→*human ancestry in 14 modern human haplotypes across the analytical parameter grid. Color indicates the proportion of parameter combinations exceeding the detectability criterion after aggregating over the remaining dimensions. Detectability is highest when introgression fractions are larger, when the ghost–Denisovan split is older than the human–Denisovan split, when pulse timing preserves multiple Denisovan-derived ancestral lineages before the ghost*→*Denisovan pulse, and when effective population sizes slow coalescence among those lineages.

Split times also changed detectability (Fig. 2b). Across split-time cells, the detectable proportion ranged from 57.6% to 93.9%. Detectability was lowest when the ghost–Denisovan split was recent relative to the human–Denisovan split, because ghost ancestry then added little extra coalescent depth. In the grid, the lowest split-time cell paired a ghost–Denisovan split of 1.0 Ma with a human–Denisovan split of 0.9 Ma. Detectability was highest when the two split times were far apart; the highest split-time cell paired a ghost–Denisovan split of 1.8 Ma with a human–Denisovan split of 0.3 Ma. The median expected shift across split-time cells ranged from 23.2 to 348.0 generations.

Effective population sizes had a measurable effect on the expected signal (Fig. 2c). Across effective-size cells, the detectable proportion ranged from 56.8% to 98.5%. The lowest value occurred when both the human and Denisovan effective population sizes were 1000, and the highest occurred when both were 10,000.

Pulse timing produced a narrower range of detectable proportions than the introgression fractions, split times, or effective population sizes (Fig. 2d). Across timing cells, the detectable proportion ranged from 80.4% to 93.3%. The highest value occurred when the Denisovan → human pulse was at 40 ka and the ghost→Denisovan pulse was at 100 ka. The lowest value occurred when the Denisovan → human pulse was at 80 ka and the ghost → Denisovan pulse was at 700 ka. This pattern reflects the number of Denisovan-side lineages retained between the two pulses: longer intervals before the ghost pulse allow more coalescence among Denisovan-derived lineages, reducing the probability of a mixed ghost and non-ghost Denisovan-side state.

### 2.3 Window-level summary statistics preserve genealogical signals of ghost introgression

Our analytical results above define the target signal in genealogical terms: Ghost → Denisovan human → ancestry should increase local coalescent depth in windows where some human haplotypes have unusually deep relationships to the Denisovan genome. We next asked whether this genealogical signal is reflected in observable summary statistics computed from such windows.

Because the simulations retain the true tree sequence, we could directly compare sequence-level summaries with local genealogical properties. Across 140,000 human–Denisovan haplotype-pair comparisons from 2,500 matched ghost-positive and ghost-negative windows, pairwise human–Denisovan sequence divergence increased with true local human–Denisovan coalescent time (Fig. 3a; Pearson (r=0.653), Spearman (*ρ* = 0.630)). This relationship was stronger when restricted to ghost-positive windows (Pearson (r=0.707), Spearman (*ρ* = 0.723)). Thus, the primary genealogical signal that defined ghost introgression, which is the local human–Denisovan coalescent depth, was reflected in human–Denisovan divergence, which is directly observable at sequence level.

**Fig. 3:**
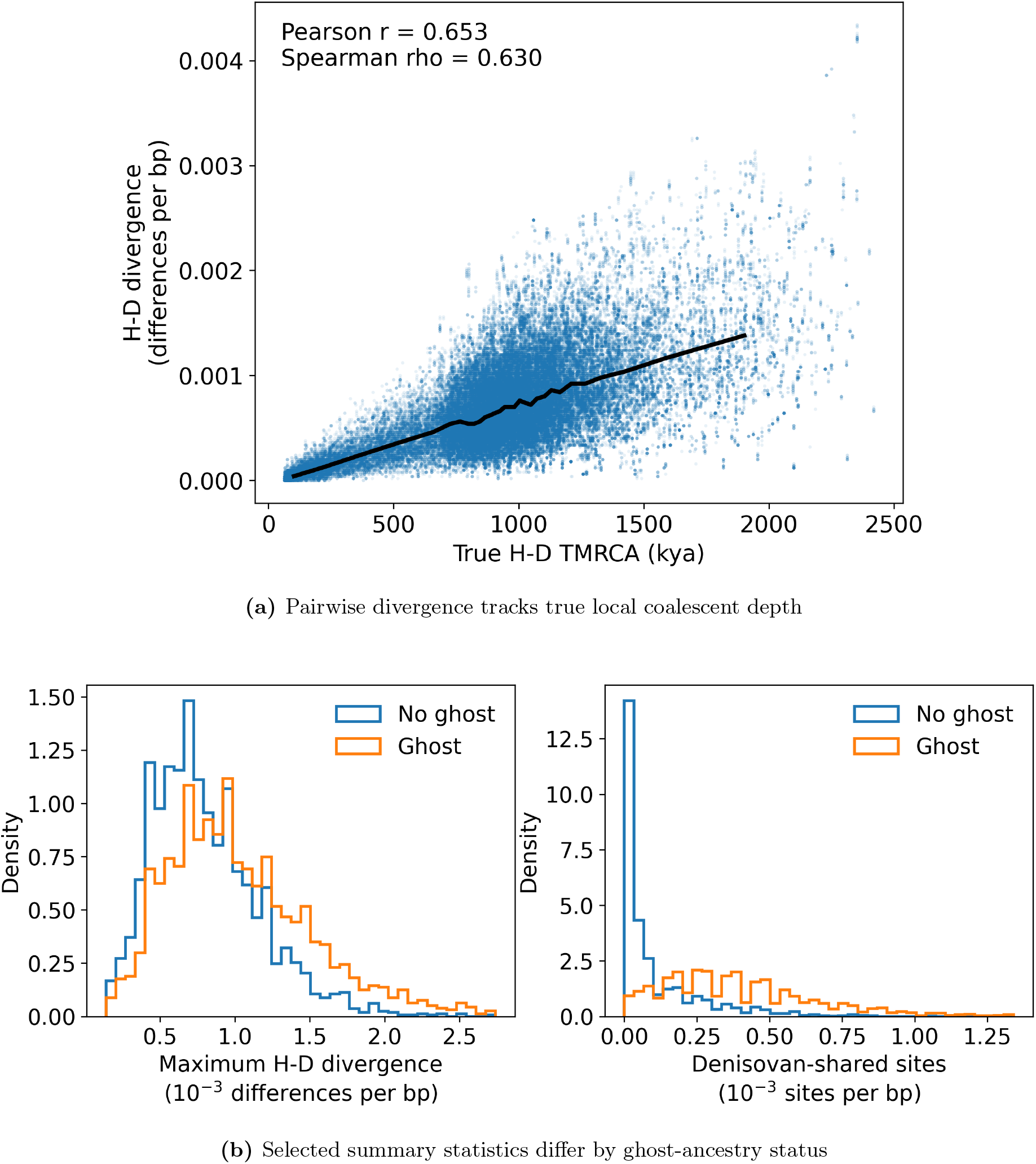
Window-level summary statistics preserve observable consequences of the genealogical signal created by Ghost*→*Denisovan*→*human ancestry. **a**, Each point represents one human–Denisovan haplotype-pair comparison in a simulated window. The black line shows the median human–Denisovan divergence within bins of true local human–Denisovan coalescent depth. **b**, Selected summary statistics differ between windows simulated with and without ghost-mediated ancestry. Maximum human–Denisovan divergence captures a tail-sensitive consequence of unusually deep local relationships, while Denisovan-shared variation captures complementary allele-sharing evidence.

Ghost-positive and ghost-negative windows also differed in window-level summaries (Fig. 3b). The clearest divergence-based signal was an increase in maximum human–Denisovan divergence, consistent with the expectation that only a subset of human haplotypes within a window may carry the ghost-mediated tract. The corresponding genealogical summaries showed the same pattern, with ghost-positive windows showing substantially higher maximum human–Denisovan coalescent times. Mean human–Denisovan divergence showed the same qualitative direction and is shown as a supporting summary in Appendix Fig. 6. Ghost positive windows also contained more Denisovan-shared variation than matched ghost-negative windows.

These analyses show that substantial simulated genealogical signal of Ghost → Denisovan → human ancestry is preserved in observable sequence summaries. Individual statistics are not sufficient classifiers on their own, but collectively retain measurable information about the genealogical distortions induced by Ghost ancestry. This result provides a link between the analytical coalescent predictions and developing a summary statistics-based classifier.

### 2.4 DEEP identifies ghost-introgressed windows and remains strongest as a high-specificity screening signal

We next trained population-specific DEEP classifiers using the summary statistics described in Methods. Simulated 50-kb windows were labeled depending on whether modern human haplotypes carried ghost ancestry. The full training workflow (Fig. 4a) required 436.0 CPU hours, most of which was spent generating labeled simulated genotype data (Table 1).

**Table 1:** Computational resources used for simulation-based classifier training. Genotype simulation and feature construction were run as 25-task arrays for each population-specific model, whereas neural-network training was run as a single final job per model. For array jobs, wall time is the maximum elapsed time among array tasks, approximating elapsed time when tasks are run concurrently; CPU-hours are summed across tasks. Peak RSS is the maximum resident set size reported by `/usr/bin/time –v`.

| Model | Stage | Wall time | CPU-hours | Peak RSS (GiB) |
| --- | --- | --- | --- | --- |
| Oceanian | Genotype simulation | 4:57:01 | 107.6 | 1.1 |
|  | Feature construction | 0:21:11 | 7.7 | 2.1 |
|  | Neural-net training | 8:56:13 | 8.3 | 15.5 |
| East Asian | Genotype simulation | 12:34:08 | 295.8 | 1.3 |
|  | Feature construction | 0:21:42 | 8.0 | 2.1 |
|  | Neural-net training | 9:19:57 | 8.7 | 16.1 |
| Total | Full workflow | – | 436.1 | 16.1 |

**Table 2:**
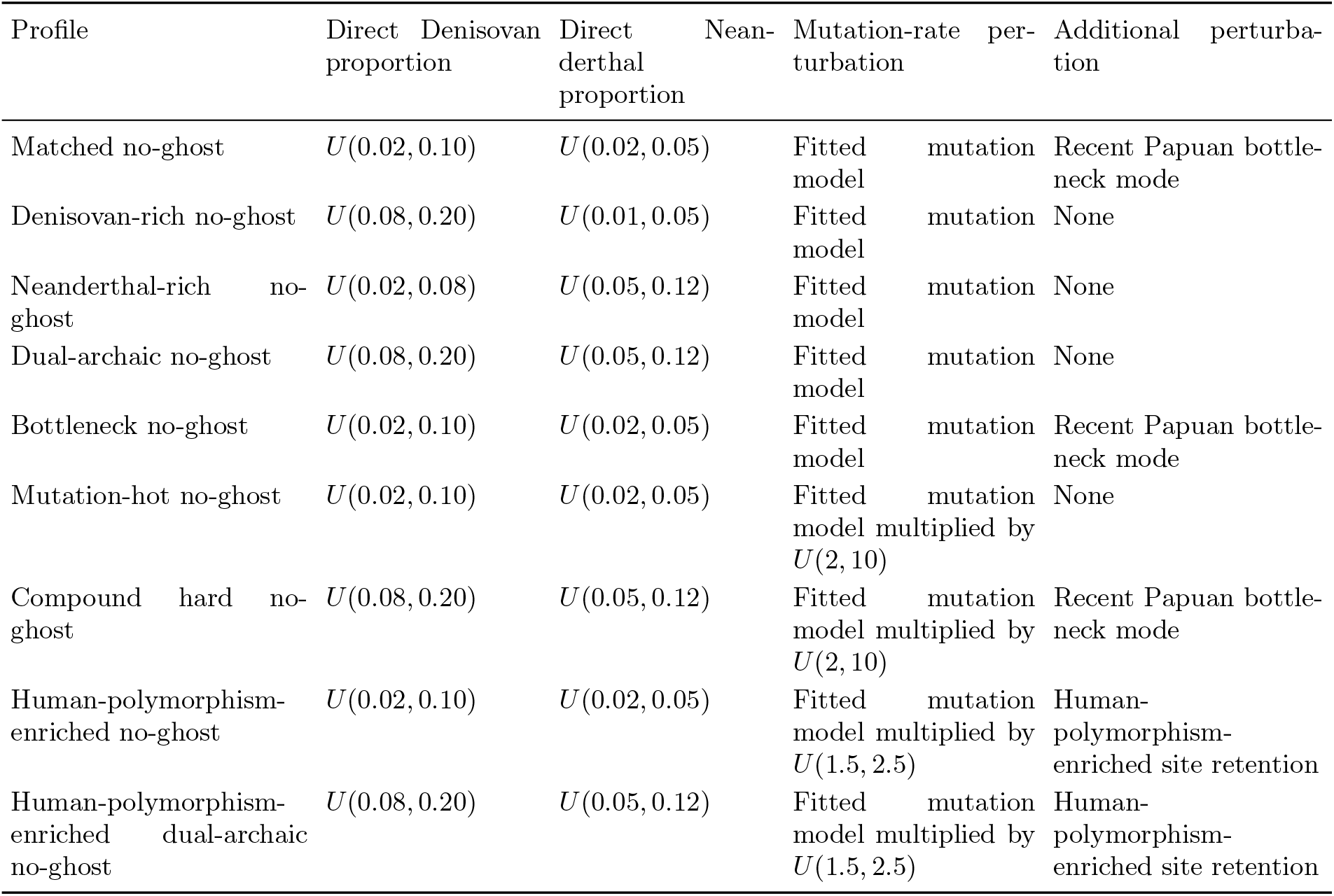
Simple Oceanian no-ghost robustness profiles. All profiles set the ghost-to-Denisovan introgression proportion to zero.

**Table 3:**
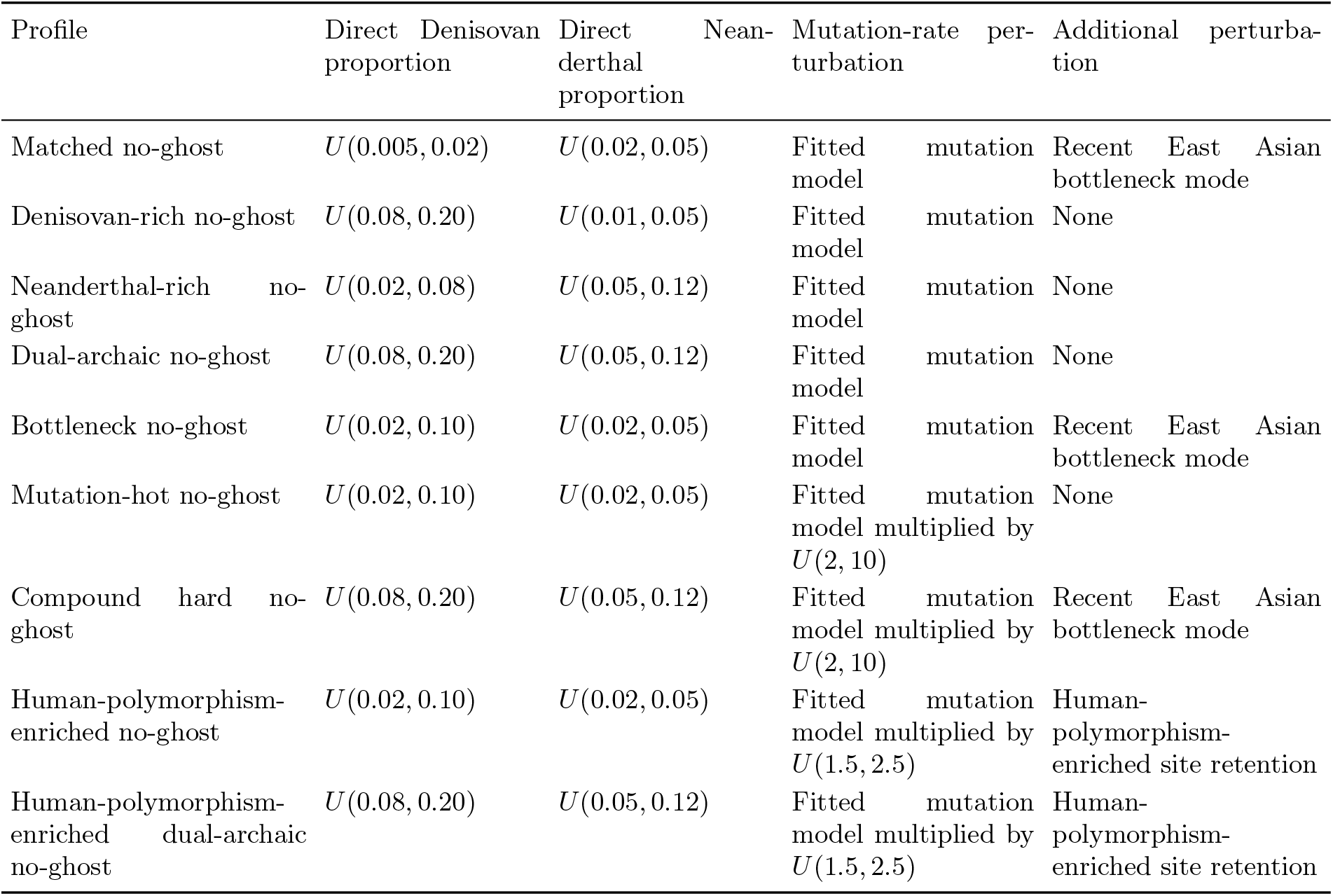
Simple East Asian no-ghost robustness profiles. All profiles set the ghost-to-Denisovan introgression proportion to zero.

**Table 4:**
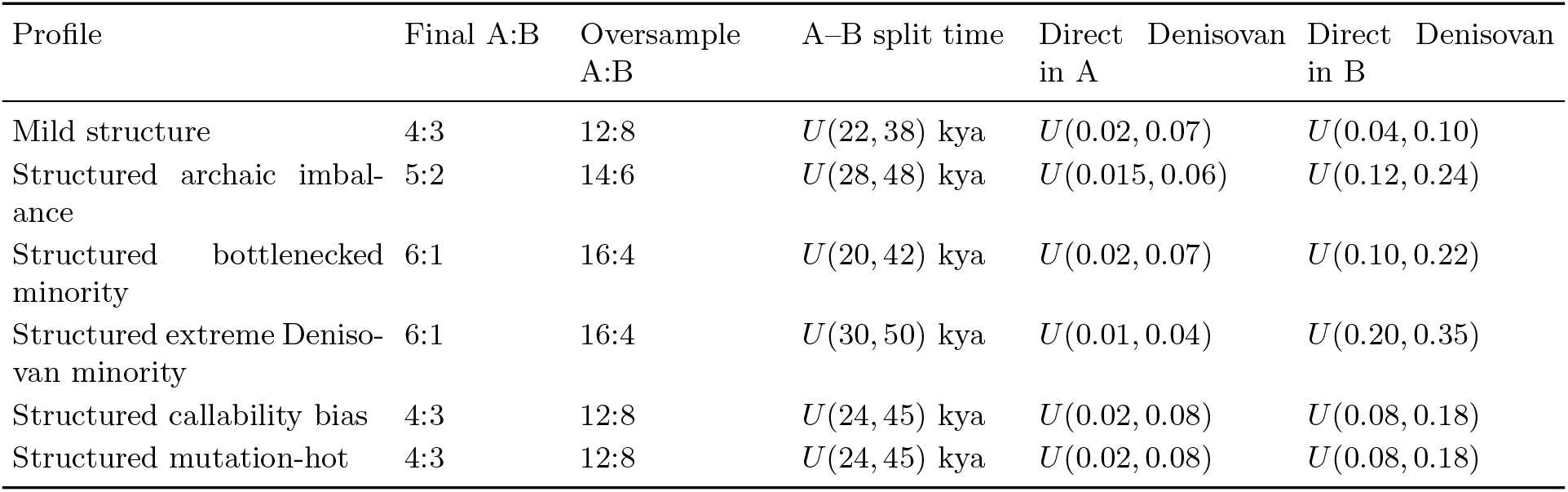
Structured sampled-human no-ghost sampling and direct Denisovan profiles.

**Table 5:**
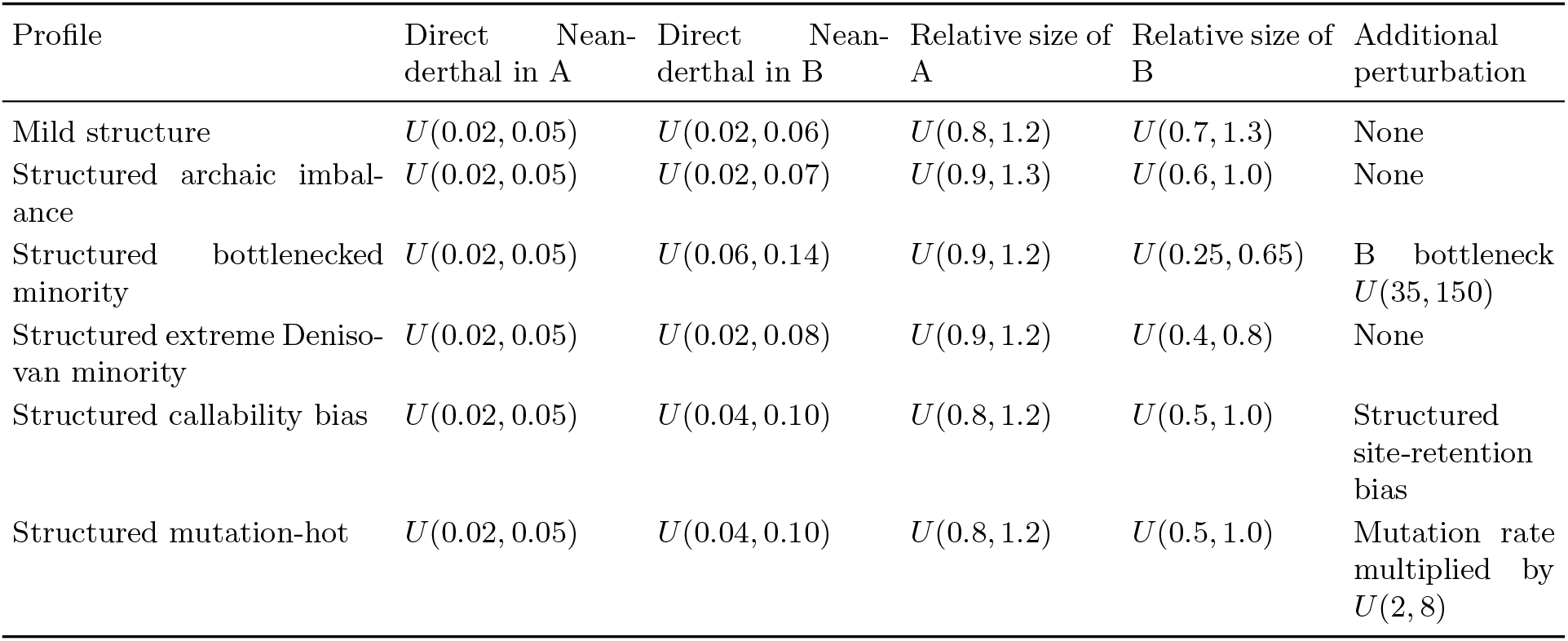
Structured sampled-human no-ghost direct Neanderthal, size, and technical perturbation profiles. Relative sizes are multipliers of the structured parent population size.

**Table 6:** Deep ancestral-structure no-ghost robustness profile. The same profile definition was used for the Oceanian and East Asian robustness analyses, with the sampled human parent population set to Papuan or East Asian, respectively.

| Parameter | Distribution or setting |
| --- | --- |
| Ghost-to-Denisovan introgression proportion | 0 |
| Direct Denisovan ancestry into sampled human parent | $U(0.03, 0.12)$ |
| Direct Neanderthal ancestry into sampled human parent | $U(0.01, 0.06)$ |
| Denisovan–Neanderthal split time | $U(350, 550)$ kya |
| AMH-side ancestral split time | $U(220, 300)$ kya |
| Entry into deep ancestral demes | $U(550, 750)$ kya, constrained to be older than the Denisovan–Neanderthal and AMH-side split times |
| Deep ancestral deme split time | $U(1200, 2200)$ kya, constrained to be older than entry into the deep ancestral demes |
| Symmetric migration rate between the deep human-side and deep archaic-side ancestors | $U(10^{-7}, 2 \times 10^{-5})$ |
| Effective size of the deep human-side ancestor | $U(2000, 12000)$ |
| Effective size of the deep archaic-side ancestor | $U(1000, 8000)$ |
| Effective size of the deep ancestral root | $U(5000, 20000)$ |
| Mutation-rate perturbation | Fitted mutation model |
| Callability perturbation | None |

**Table 7:** Robustness of the Oceanian and East Asian classifiers across no-ghost perturbation scenarios, including direct archaic ancestry, sampled-human structure, deep ancestral structure, ascertainment bias, and mutation-rate elevation.

| Scenario | Median | 95th | 99th | FPR $\leq 1/55,000$ | FPR $\leq 1/1,000$ |
| --- | --- | --- | --- | --- | --- |
| Matched no-ghost | 0.099 | 0.808 | 0.909 | 0 (0.000%) | 0 (0.000%) |
| Denisovan-rich | 0.603 | 0.856 | 0.921 | 0 (0.000%) | 0 (0.000%) |
| Dual archaic | 0.589 | 0.847 | 0.955 | 0 (0.000%) | 2 (0.400%) |
| Mild structure | 0.590 | 0.854 | 0.919 | 0 (0.000%) | 0 (0.000%) |
| Structured archaic imbalance | 0.631 | 0.825 | 0.916 | 0 (0.000%) | 0 (0.000%) |
| Structured bottlenecked minority | 0.584 | 0.816 | 0.894 | 0 (0.000%) | 1 (0.200%) |
| Deep ancestral structure | 0.600 | 0.969 | 0.985 | 0 (0.000%) | 13 (2.600%) |
| Human-poly enriched | 0.401 | 0.918 | 0.957 | 0 (0.000%) | 4 (0.800%) |
| Mutation-hot | 0.946 | 1.000 | 1.000 | 85 (17.000%) | 207 (41.400%) |

East Asian
| Scenario | Median | 95th | 99th | FPR $\leq 1/55,000$ | FPR $\leq 1/1,000$ |
| --- | --- | --- | --- | --- | --- |
| Matched no-ghost | 0.076 | 0.669 | 0.884 | 0 (0.000%) | 0 (0.000%) |
| Denisovan-rich | 0.530 | 0.805 | 0.904 | 0 (0.000%) | 1 (0.200%) |
| Dual archaic | 0.510 | 0.809 | 0.892 | 0 (0.000%) | 1 (0.200%) |
| Mild structure | 0.482 | 0.762 | 0.854 | 0 (0.000%) | 0 (0.000%) |
| Structured archaic imbalance | 0.515 | 0.794 | 0.889 | 0 (0.000%) | 0 (0.000%) |
| Structured bottlenecked minority | 0.468 | 0.779 | 0.892 | 0 (0.000%) | 2 (0.400%) |
| Deep ancestral structure | 0.635 | 0.968 | 0.994 | 2 (0.400%) | 23 (4.600%) |
| Human-poly enriched | 0.629 | 0.942 | 0.990 | 0 (0.000%) | 11 (2.200%) |
| Mutation-hot | 0.995 | 1.000 | 1.000 | 211 (42.200%) | 321 (64.200%) |

All windows were simulated without Ghost → Denisovan ancestry. The Median, 95th, and 99th columns report summaries of the classifier score distribution for each no-ghost scenario. Entries in the final two columns give the number of windows exceeding the ROC-calibrated threshold, with the corresponding percentage in parentheses. The deep ancestral-structure profile tests whether ancestral population structure across the hominin tree can inflate ghost-ancestry scores in the absence of Ghost → Denisovan introgression. Complete scenario descriptions, sample counts, and additional fixed-score thresholds are provided in the machine-readable supplementary CSV.

**Fig. 4:**
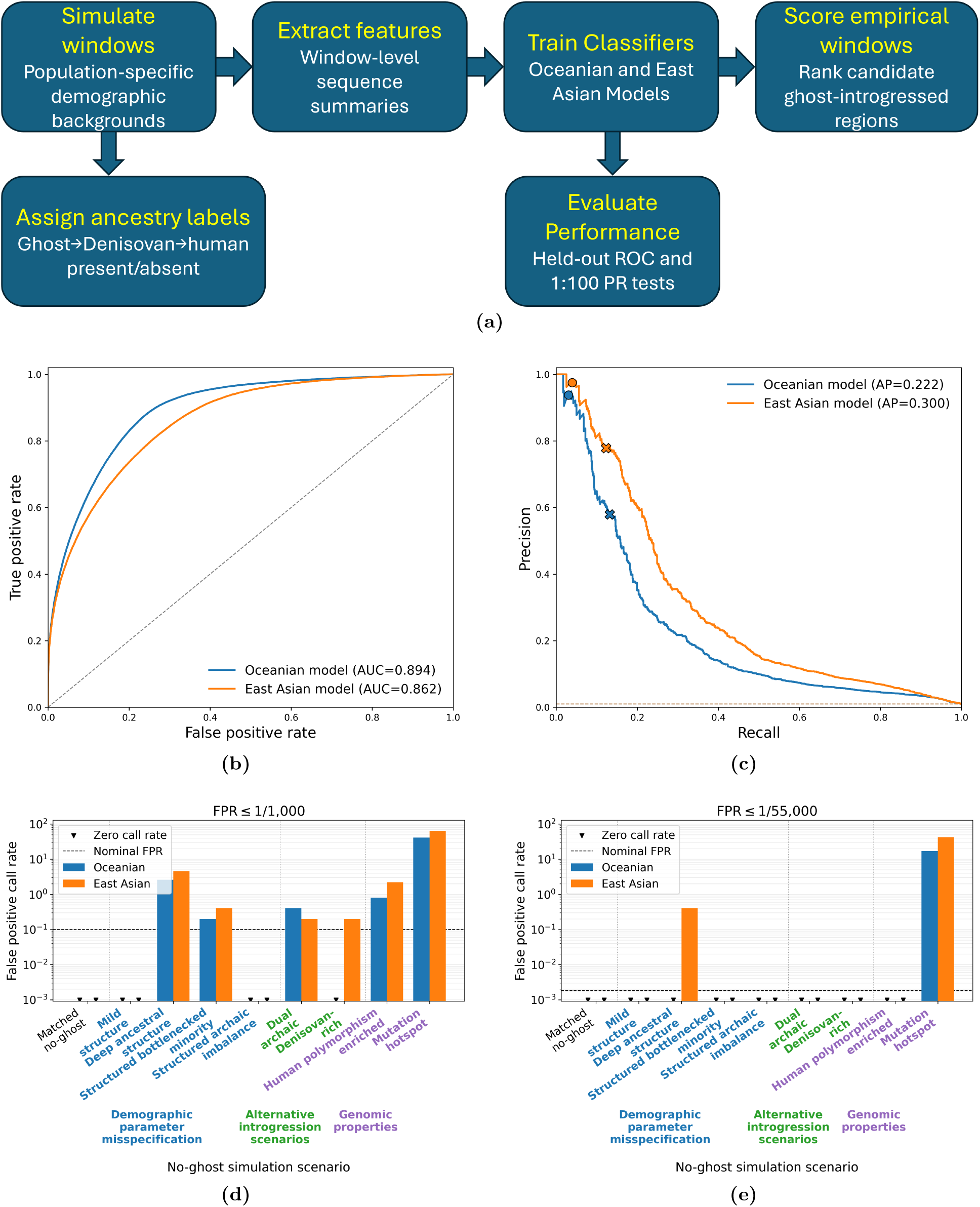
DEEP workflow, held-out performance, and robustness analysis. **a**, DEEP workflow for identifying candidate Ghost*→*Denisovan*→*human windows. Simulations generate genomic windows with known ancestry labels. Observable window-level summary statistics are extracted from simulated genotype data and used to train population-specific neural-network classifiers. The trained models are then evaluated on held-out simulations and applied to empirical windows to produce candidate scores for ghost-mediated ancestry. **b**, ROC curves computed on balanced held-out test sets. **c**, Precision–recall curves computed on 1:100 skewed evaluation sets. Points show the performance obtained by applying thresholds selected from the ROC analysis. Circles mark the threshold maximizing TPR subject to FPR ≤ 1/55,000, and X marks indicate the threshold maximizing TPR subject to FPR ≤ 1/1,000. These thresholds are projected onto the precision–recall curves to show their precision and recall under class imbalance. **d**, Robustness analysis at the FPR ≤ 1/1,000 threshold. Bars show the false-positive call rate in no-ghost simulation scenarios, grouped by demographic-parameter misspecification, alternative introgression scenarios, and genomic properties. The dashed line marks the nominal FPR used to calibrate the threshold. **e**, Robustness analysis at the stricter FPR ≤ 1/55,000 threshold, using the same no-ghost simulation scenarios and grouping scheme as in panel **d**. Zero-call scenarios are shown at the plotting floor because the y-axis is log-scaled.

Both the East Asian and Oceanian models showed strong overall discrimination between ghost-positive and ghost-negative windows (Fig. 4b), achieving ROC AUC of 0.894 and 0.862 respectively for Oceanians and East Asians. These results indicate that the window-level features learned by the networks captured reproducible signals associated with superarchaic ancestry.

For genome-wide scans, however, performance at very low false positive rates is more relevant than average discrimination across all thresholds. We therefore identified score cutoffs that maximized true positive rates (TPRs) while constraining false positive rates (FPRs) to either FPR≤1/55,000, corresponding to one expected false positive across a genome-wide scan, or FPR ≤ 1/1,000, a less stringent but still conservative screening threshold. At FPR ≤ 1/55,000, the Oceanian and East Asian classifiers achieved TPRs of 0.023 and 0.044 respectively. Relaxing the threshold to FPR ≤ 1/1,000 increased TPRs to 0.120 and 0.123 respectively.

We then evaluated precision–recall performance under a more realistic 1:100 class imbalance (Fig. 4c). Average precision reached 0.222 for the Oceanian model and 0.300 for the East Asian model, substantially exceeding the 1% positive-class prevalence. As expected, the choice of threshold preduced a tradeoff between precision and recall. At FPR ≤ 1/55,000, both Oceanian and East Asian models produced high precision (0.938 and 0.975) but low recall (0.030 and 0.039. Relaxing the threshold to FPR ≤ 1/1,000 increased recall (0.132 and 0.123) but at the cost of reduced precision (0.579 and 0.778).

Together, these results show that DEEP is most effective as a high-specificity screening framework. At a stricter FPR ≤ 1/55,000 threshold, precision was high but recall was low, whereas at the more relaxed FPR ≤ 1/1,000 threshold, recall increased nearly four-fold with lower precision. We therefore carried both thresholds forward, using one for highest-specificity calls and the other for broader analyses. We evaluated these calibrated operating points in subsequent robustness analyses to test whether plausible demographic or genomic misspecification inflates false-positive calls (Fig. 4d and 4e).

### 2.5 Conservative thresholds are robust to alternative demographic and genomic scenarios

We next evaluated whether DEEP assigns high ghost-ancestry scores to simulated windows that contain no ghost ancestry but depart from the training models in ways that could plausibly mimic the target signal. In all robustness simulations, the Ghost → Denisovan pulse was removed, such that any high-scoring windows represent false-positive calls. We considered three classes of perturbations: demographic misspecifications, alternative introgression histories, and genomic-property perturbations (Fig. 4d and 4e).

Alternative-introgression scenarios produced low false-positive rates (Fig. 4d and 4e). In the Oceanian classifier, the Denisovan-rich scenario produced no calls at either the strict 1/55,000 threshold or the relaxed 1/1,000 threshold, despite a median score of 0.603 and a 99th percentile score of 0.921. The dual-archaic scenario likewise produced no calls at the strict threshold and only 2 of 500 calls at the relaxed threshold, corresponding to a 0.4% false-positive rate. The East Asian classifier showed the same pattern: the Denisovan-rich and dual-archaic scenarios produced no strict-threshold calls and only 1 of 500 relaxed-threshold call in each scenario, corresponding to 0.2% false-positive rates. Thus, increasing direct Denisovan ancestry or adding Neanderthal ancestry did not by itself produce appreciable numbers of high-scoring windows under either operating threshold.

Most demographic parameter misspecifications also produced low call rates. For the Oceanian classifier, mild sampled-human structure, archaic-ancestry imbalance between populations, and a bottlenecked-minority population produced no strict-threshold calls, and relaxed-threshold call rates of 0%, 0%, and 0.2%, respectively. Their 99th percentile scores ranged from 0.894 to 0.919. For the East Asian classifier, the same three scenarios again produced no strict-threshold calls, and relaxed-threshold call rates of 0%, 0%, and 0.4%, respectively, with 99th percentile scores ranging from 0.854 to 0.892. These results indicate that the classifier scores were not strongly inflated by the sampled-human structure and population-size perturbations considered here.

The deep ancestral-structure scenario represented the most stringent biological stress test. In this scenario, modern and archaic humans descend through distinct ancestral populations connected by low migration prior to their common ancestry, generating unusually deep coalescent relationships without ghost introgression. This setting increased score distributions in both classifiers. In the Oceanian classifier, the median, 95th percentile, and 99th percentile scores were 0.600, 0.969, and 0.985, respectively; the relaxed threshold called 13 of 500 windows, for a 2.6% false-positive rate, while the strict threshold called none. In the East Asian classifier, the corresponding score summaries were 0.635, 0.968, and 0.994; the relaxed threshold called 23 of 500 windows, for a 4.6% false-positive rate, while the strict threshold called 2 of 500 windows, or 0.4%. Thus, deep ancestral structure increased DEEP scores relative to other demographic perturbations, but most windows remained below threshold, especially under the stricter operating point.

The largest false-positive inflation occurred under mutation-rate elevation (Fig. 4d and 4e). This scenario was intentionally severe: the simulated mutation rate was centered sixfold above the corresponding fitted genome-wide mutation-rate model, approximately 6.0× 10^−8^for the Oceanian/Papuan model and 6.6 ×10^−8^ for the East Asian model, rather than the fitted means of 1.0 ×10^−8^ and 1.1 ×10^−8^, respectively. Under this extreme perturbation, the Oceanian classifier had a median score of 0.946 and produced 85 of 500 calls at the strict threshold and 207 of 500 calls at the relaxed threshold, corresponding to false-positive rates of 17.0% and 41.4%. The East Asian classifier was still more sensitive, with a median score of 0.995 and false-positive rates of 42.2% at the strict threshold and 64.2% at the relaxed threshold. In contrast, human-polymorphism enrichment produced much smaller inflation: no calls at the strict threshold in either classifier, and relaxed-threshold false-positive rates of 0.8% in the Oceanian classifier and 2.2% in the East Asian classifier. The matched no-ghost baseline produced no calls in either classifier at either threshold.

Together, these no-ghost robustness tests indicate that conservative DEEP calls are not substantially inflated by additional archaic ancestry or most demographic misspecifications, while also identifying extreme mutation-rate elevation as the main non-ghost condition capable of increasing classifier scores.

### 2.6 Empirical superarchaic-ancestry scores identify sparse, shared, and externally enriched candidate regions

We next applied DEEP to empirical 50-kb windows from Oceanians in the Simons Genome Diversity Project [46], Tibetans from Lu et al. [47], and Han Chinese from the high-coverage 1000 Genomes Project/IGSR genomes [48–50] to summarize the genomic distribution of candidate Denisovan-mediated superarchaic ancestry. The empirical datasets contained 51,726 Oceanian windows, 52,771 Tibetan windows, and 52,704 Han Chinese windows. The calibrated FPR≤ 1/1,000 thresholds were 0.9760 for the Oceanian model and 0.9733 for the East Asian model, with observed held-out false-positive rates of 9.97 ×10^−4^ and 9.92× 10^−4^, respectively. The stricter FPR ≤ 1/55,000 thresholds were 0.9998 for the Oceanian model and 0.998 for the East Asian model, with observed held-out false-positive rates of 1.60× 10^−5^ for both models. Using the FPR≤ 1/1,000 threshold, high-scoring windows comprised 130/51,726 windows in Oceanians (0.25%), 323/52,771 windows in Tibetans (0.61%), and 231/52,704 windows in Han Chinese (0.44%). At the FPR ≤ 1/55,000 threshold, the corresponding counts were 17/51,726 windows in Oceanians (0.03%), 86/52,771 windows in Tibetans (0.16%), and 27/52,704 windows in Han Chinese (0.05%).

Predicted scores were positively correlated across populations. Across 51,609 windows matched in all three empirical datasets, the strongest pairwise correlation was observed between Tibetans and Han Chinese with Pearson r = 0.458 and Spearman *ρ* = 0.426. Oceanians and Tibetans had Pearson r = 0.429 and Spearman *ρ* = 0.419, and Oceanians and Han Chinese had Pearson r = 0.343 and Spearman *ρ* = 0.340 (Fig. 5c). We also quantified overlap among thresholded DEEP-positive sets. Within the matched-window set, at the FPR≤ 1/1,000 threshold, 62 windows were shared between Oceanians and Tibetans, 34 between Oceanians and Han Chinese, and 75 between Tibetans and Han Chinese. These corresponded to Jaccard indices of 0.161, 0.104, and 0.158, respectively. At the FPR≤ 1/55,000 threshold, 13 windows were shared between Oceanians and Tibetans, 7 between Oceanians and Han Chinese, and 18 between Tibetans and Han Chinese, corresponding to Jaccard indices of 0.144, 0.189, and 0.189. These overlaps were higher than expected under random overlap of high-scoring windows. Across all matched windows, observed overlap at the FPR ≤ 1/1,000 threshold was 38.3–52.7 times higher than the chromosome-stratified expectation, with Fisher exact-test odds ratios ranging from 92.2 to 182.4 across population pairs (all Fisher *P ≤* 1.64 *×* 10^−50^; all chromosome-stratified permutation *P ≤* 1.0 *×* 10^−5^). At the stricter FPR ≤ 1/55,000 threshold, observed overlap was 143.5–164.7 times higher than the chromosome-stratified expectation, with Fisher exact-test odds ratios ranging from 1515.1 to 2293.7 (all Fisher *P ≤* 8.90 *×*10^−20^; all chromosome-stratified permutation *P ≤* 1.0 *×* 10^−5^).

**Fig. 5:**
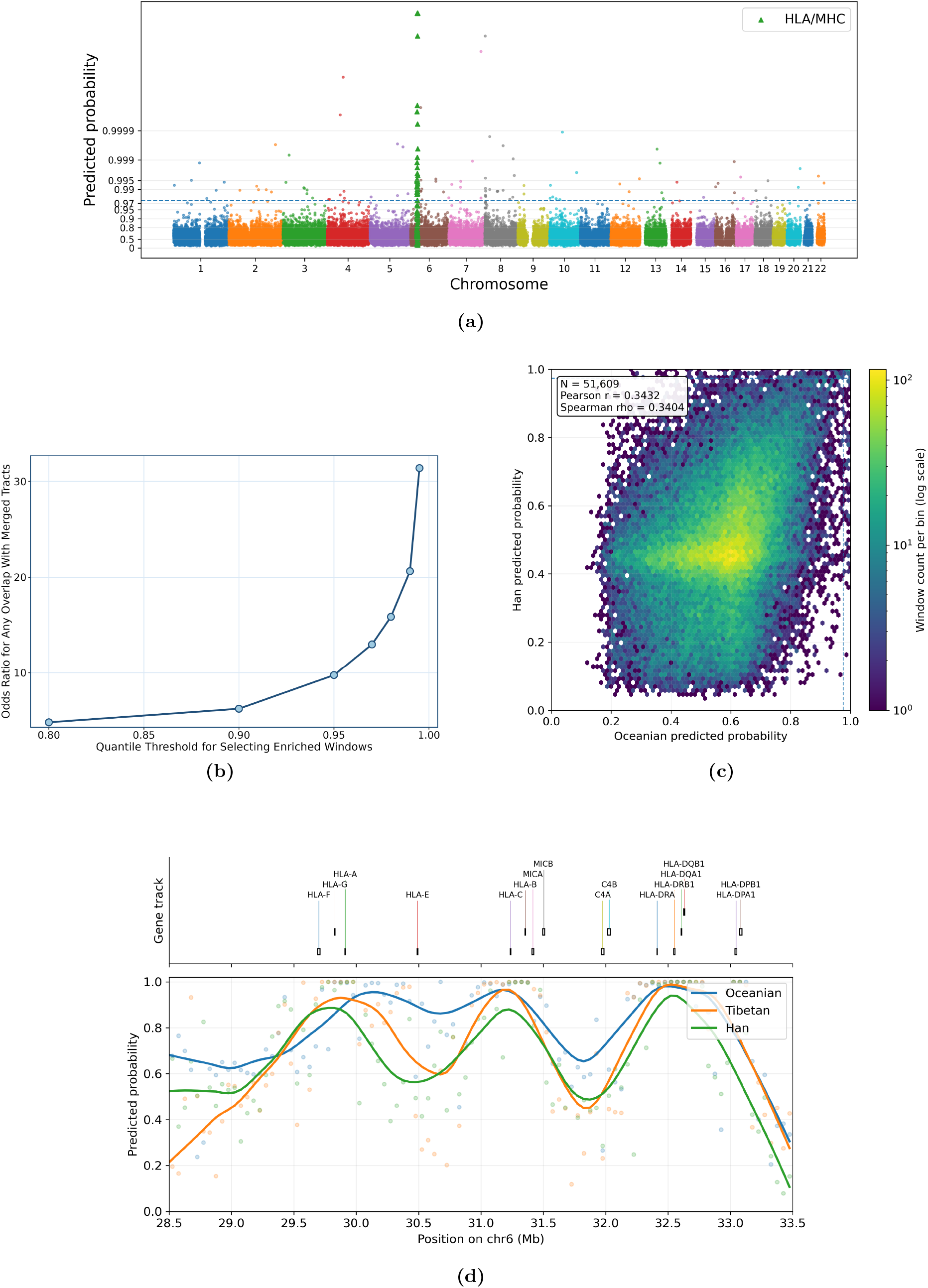
Empirical superarchaic-ancestry score landscape, external enrichment, and HLA/MHC locus-level profiles. **a**, Manhattan plot of Oceanian predicted probabilities across autosomal 50 kb windows. The horizontal line marks the FPR ≤ 1/1,000 threshold, and the probability axis is expanded near one to make extreme high-scoring windows visually distinguishable. **b**, Enrichment of high-scoring Oceanian windows in Papuan-private variant tracts. Points show odds ratios comparing DEEP-positive and DEEP-negative windows for overlap with tracts constructed from windows enriched for variants present in Papuans and absent from Africans across top-quantile thresholds. **c**, Matched-window comparison of Oceanian and Han predicted probabilities. Hexagonal bins show the density of empirical windows, and dashed lines mark the corresponding FPR ≤ 1/1,000 thresholds. **d**, Zoom-in across the GRCh37/hg19 HLA/MHC interval chr6:28.5–33.5 Mb. The upper track shows representative HLA/MHC genes, and the lower panel shows smoothed predicted-probability profiles for Oceanians, Tibetans, and Han.

**Fig. 6:**
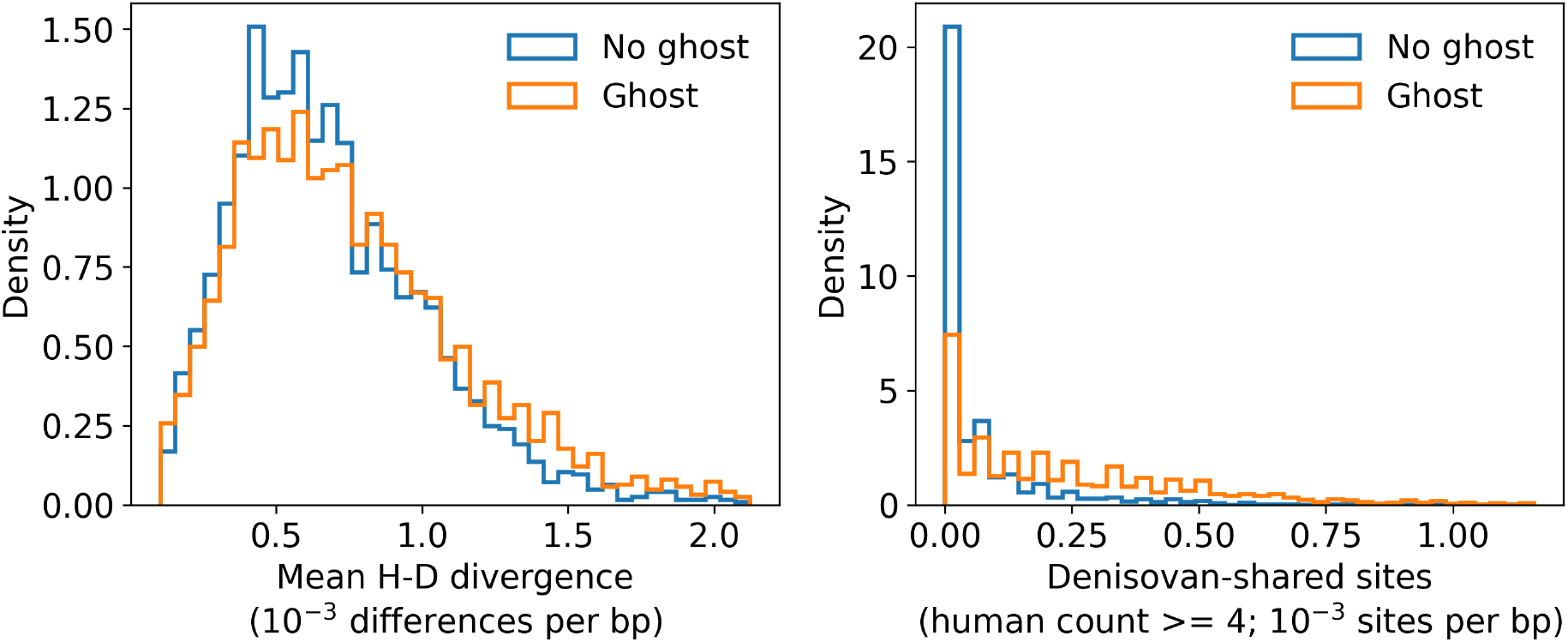
Additional summary-statistic shifts in simulated ghost-positive and ghost-negative windows. Mean human– Denisovan divergence and stricter Denisovan-shared variation, requiring the alternate allele to be present in at least four human haplotypes, show the same qualitative direction as the main summaries in Fig. 3. These panels are presented as supporting checks rather than as the primary visual evidence for the summary-statistic signal.

**Fig. 7:**
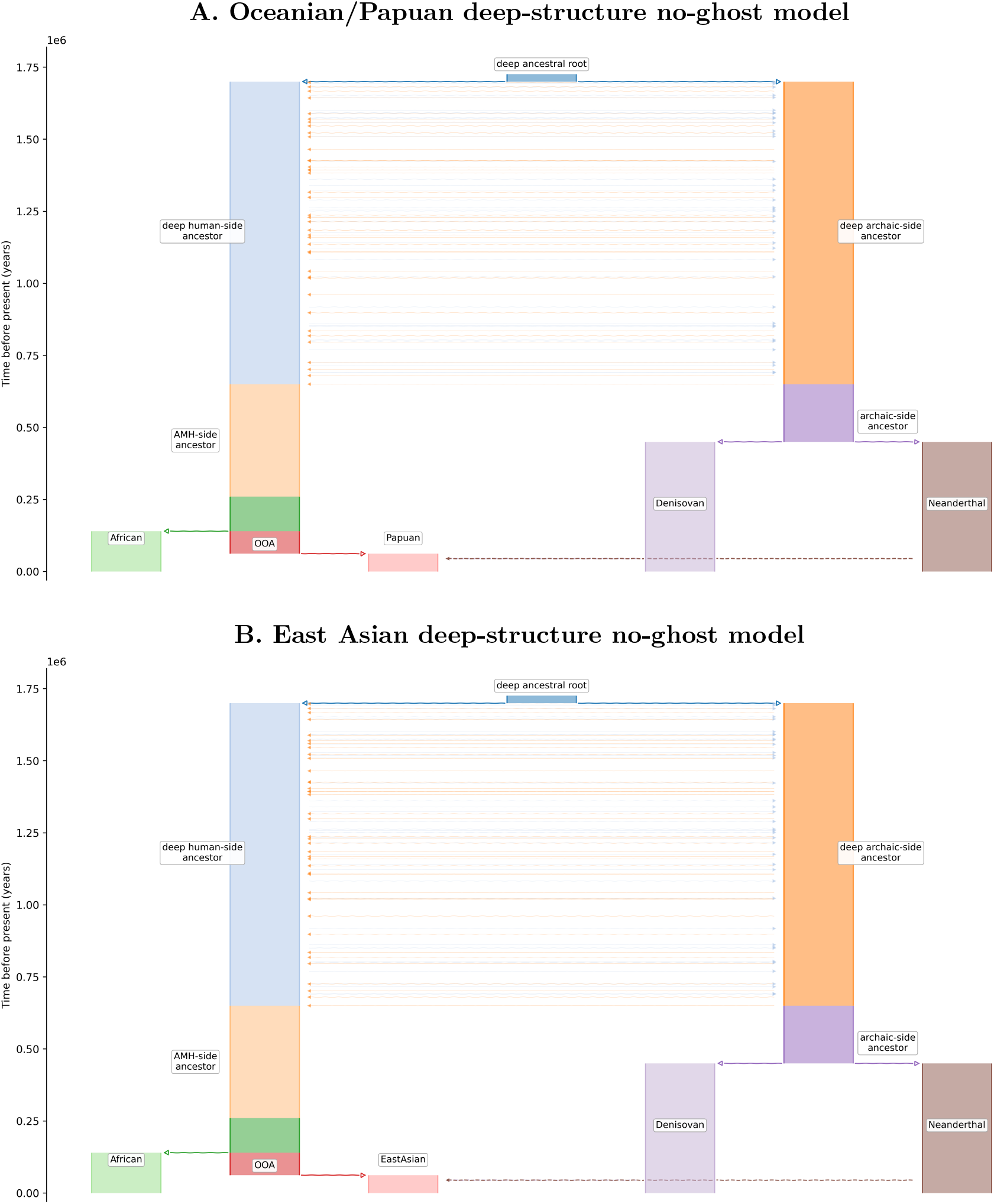
Deep ancestral-structure no-ghost robustness models. Each panel shows the corresponding demography used to test whether ancestral population structure across the hominin tree can generate high ghost-ancestry scores in the absence of Ghost*→*Denisovan introgression. Both versions include a deep human-side ancestral deme, a deep archaic-side ancestral deme, low migration between the two deep demes, and direct Denisovan and Neanderthal ancestry into the sampled human parent. The Ghost*→*Denisovan pulse was set to zero in all replicates.

**Fig. 8:**
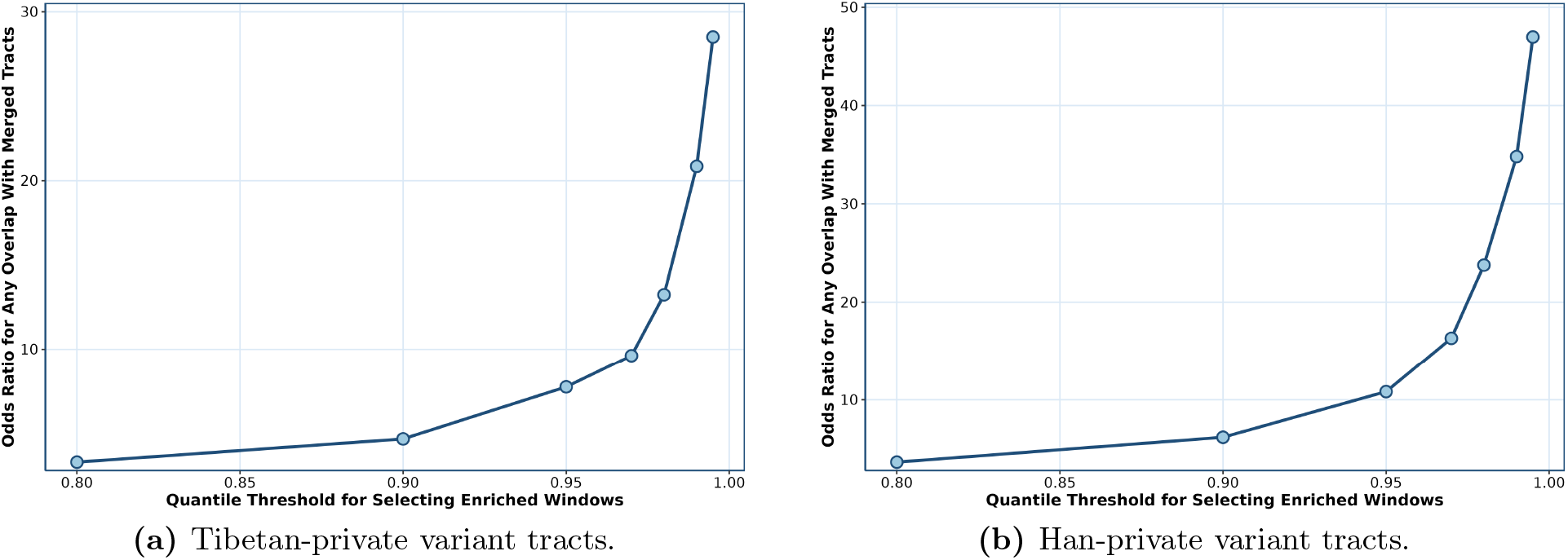
Enrichment of high-scoring East Asian windows in population-private variant tracts. Points show odds ratios comparing DEEP-positive and DEEP-negative 50 kb windows for overlap with merged tracts constructed from windows enriched for variants present in the focal population and absent from Africans across top-quantile thresholds.

Chromosome 6 contained the largest number of DEEP-positive windows in all three populations. At the FPR ≤ 1/1,000 threshold, chromosome 6 contained 42 DEEP-positive windows in Oceanians, 46 in Tibetans, and 34 in Han Chinese, corresponding to 1.28%, 1.38%, and 1.02% of analyzed chromosome-6 windows, respectively. At the stricter FPR ≤ 1/55,000 threshold, chromosome 6 contained 11 DEEP-positive windows in Oceanians, 23 in Tibetans, and 13 in Han Chinese. These values corresponded to 0.33%, 0.69%, and 0.39% of analyzed chromosome-6 windows. Although high-scoring windows also occurred on other chromosomes, the repeated chromosome-6 signal motivated a closer examination.

The chromosome 6 signal was concentrated in the HLA/MHC interval chr6:28.5–33.5 Mb (GRCh37/hg19). This interval contained 101 overlapping 50-kb windows in each population. The maximum predicted score in the interval was 1.000 in all three populations, and the mean predicted scores were 0.771 in Oceanians, 0.697 in Tibetans, and 0.666 in Han Chinese. In the smoothed locus-level profiles, the peak predicted probability was 0.980 in Oceanians near 32.475 Mb, 0.988 in Tibetans near 32.525 Mb, and 0.940 in Han Chinese near 32.525 Mb (Fig. 5d). The repeated elevation of scores in HLA/MHC is consistent with previous evidence that immune-related loci retain archaic variants [51–53]. At the same time, HLA/MHC is among the most polymorphic regions of the human genome and has been shaped by pathogen-mediated and balancing selection, both of which can preserve deeply divergent haplotypes and generate unusually deep local genealogies [54–58]. We therefore report HLA/MHC as a recurrent high-scoring region while treating the specific evolutionary mechanism at this locus as unresolved. The population pairwise overlap enrichment was not explained solely by the HLA/MHC interval: after excluding HLA/MHC windows, all three population pairs remained significantly enriched at the FPR ≤ 1/1,000 threshold (all Fisher *P≤* 1.15 × 10^−24^; all chromosome-stratified permutation *P ≤* 1.0 × 10^−5^). At the stricter FPR ≤ 1/55,000 threshold, overlap outside HLA/MHC remained significant for Oceanian–Tibetan and Tibetan– Han comparisons, whereas Oceanian–Han had no shared DEEP-positive windows outside HLA/MHC.

To evaluate whether high-scoring windows overlap independent annotations of divergent ancestry, we compared DEEP predictions with Papuan-private variant tracts, SPrime calls, and hmmix calls. In Oceanians, DEEP-positive windows were enriched for Papuan-private variant tracts, with odds ratios increasing as the private-variant definition became more stringent and exceeding twenty-fold at the most extreme thresholds (Fig. 5b). DEEP-positive Oceanian windows were also enriched for SPrime archaic ancestry tracts [20, 59], with an odds ratio of 6.47 by Fisher’s exact test (*P <* 2.2 ×10^−16^), and for hmmix archaic ancestry tracts [21, 60], with an odds ratio of 3.45 (*P* = 7.54 ×10^−12^). Tibetan and Han Chinese analyses also showed significant enrichment for archaic ancestry tracts defined by both SPrime and hmmix (Supplementary Material).

Together, the empirical analyses identified a sparse set of high-scoring windows across Asian and Oceanian populations. At the FPR≤ 1/1,000 threshold, DEEP-positive windows accounted for 0.25–0.61% of analyzed windows across populations; at the FPR ≤ 1/55,000 threshold, they accounted for 0.03–0.16%. Pairwise score correlations ranged from *r* = 0.343 to *r* = 0.458, and thresholded overlap was strongest between Tibetans and Han Chinese at the strict threshold by binary correlation (*r* = 0.373). The strongest recurrent locus-level signal was observed at HLA/MHC, and DEEP-positive windows in all three analyzed populations showed significant enrichment for three independent divergent-ancestry annotations.

## 3 Discussion

Despite the growing evidence of ghost and superarchaic introgression, substantial uncertainty remains regarding the magnitude, genomic distribution, and interpretation of these signals. Much of the uncertainty arises due to the expected signal being genealogical, while the donor genome is unsampled and alternative non-introgression processes can generate similar unusually deep local coalescence. The goal of this work is not only to identify candidate regions of Denisovan-mediated superarchaic ancestry, but also ask how much genealogical information can be recovered without explicit reconstruction of local genealogy.

Here we address this question by developing DEEP, an ARG-free neural-network framework for detecting candidate regions of deeply divergent ancestry through efficient window-level proxies of local genealogical structure. A central result of this study is that substantial information regarding superarchaic ancestry is preserved in compressed, observable sequence summaries. Our analytical and simulation results show that Ghost→ Denisovan →human ancestry produces predictable shifts toward deeper Human-Denisovan coalescent times, and this shift leaves measurable traces in pairwise divergence, Denisovan-shared variation, and related haplotype summaries. Applying DEEP to empirical genomes reveals a sparse but reproducible landscape of superarchaic ancestry across Oceanians, Tibetans and Han Chinese, including both shared and population-specific regions.

A central advantage of DEEP is that it provides a scalable framework without requiring genome-wide ARG reconstruction. ARG-based approaches remain powerful for reconstructing evolutionary history, but they are computationally intensive, sensitive to modeling assumptions, and difficult to deploy repeatedly across many populations and demographic scenarios. In contrast, once a population-specific demographic model has been specified and a classifier is trained, DEEP can be applied genome-wide by computing window-level summary statistics without reconstructing local genealogies. This scalability is complementary to another practical advantage, which is, the empirical analyses presented here used only seven diploid genomes per population together with one single Denisovan genome. This makes the approach potentially useful for understudied populations where very large cohorts are not available, provided that the relevant ascertainment, filtering, and demographic assumptions can be modeled carefully.

Our results broadly converged with several recent studies suggesting that Denisovans carried superarchaic ancestry that later entered modern human populations, despite the fundamental methodological differences. At conservative thresholds, DEEP identified only a small fraction of autosomal windows as candidates for superarchaic ancestry. At the same time, these candidate regions were not randomly distributed. Scores were positively correlated across populations, high-scoring windows overlapped more often than expected by chance, and calls were enriched for population-private variant tracts as well as archaic ancestry tracts independently inferred by SPrime and hmmix.

Notably, the empirical landscape was neither completely shared nor entirely population-specific. Several biological processes could contribute to this pattern. For example, Denisovan populations themselves were likely heterogenous, and different modern populations may have inherited superarchaic ancestry from distinct Denisovan groups. Furthermore, population-specific selection on deleterious introgressed variants, drift, recombination, demography, and the retention of beneficial alleles in different ecological contexts may have all modified the original signal. Additionally, DEEP operates in a high-specificity but relatively low-recall regime, incomplete overlap among high-scoring windows should be interpreted cautiously, as truly shared regions may pass the significance thresholds in some populations but not others.

The repeated high scores near HLA/MHC require especially careful interpretation. Superarchaic ancestry being present in this region is biologically plausible because immune-related loci are repeatedly reported targets for adaptive introgression. However, HLA/MHC is also one of the clearest examples of long-term balancing selection, which also preserves deeply diverged haplotypes without recent introgression. Because DEEP is trained to detect unusually deep coalescent signal, HLA/MHC is precisely the kind of region where superarchaic ancestry, ordinary archaic introgression, balancing selection, and unusual haplotype structure may be difficult to disentangle. We therefore interpret the HLA/MHC signal cautiously. It should not be treated as definitive evidence for superarchaic ancestry by itself. Nonetheless, its repeated elevation across all three populations is notable and suggests that immune-related regions may be especially informative targets for future local haplotype, ARG-based, and functional analyses.

Our results also intersect with the recent paleoproteomic evidence suggesting that some Denisovan superarchaic ancestry may ultimately derive from populations related to *H. erectus* [17, 18, 61, 62]. Although such interpretations remain tentative given the limited information from ancient proteins relative to genomes, these findings illustrate that genomics, fossils and paleoproteomics can converge into a unified picture of understanding the biological identities of currently unsampled hominin lineages. However, it should be treated with caution as the relationship between those protein variants and the genomic signals detected in Denisovans and modern humans remains uncertain. Even so, the multiple lines of evidence point toward a more complex history of hominin interaction in Asia than can be reconstructed from the currently available ancient DNA samples alone.

There are several aspects of out analysis that need to be considered when evaluating our results. First, the classifier does not infer the identity of the ghost donor, the exact timing of the ghost-to-Denisovan event, or the full demographic history that generated each empirical window. The model asks whether empirical windows resemble simulations containing Ghost → Denisovan →human ancestry under specified population-specific backgrounds. A high score is therefore evidence of similarity to the simulated target class, not a direct observation of donor ancestry. Second, we did not explicitly test the causal mechanism responsible for differences among populations. Shared and population-specific calls may reflect differences in Denisovan donor ancestry, selection, drift, local mutation rate, local ancestry retention, or power, and these alternatives require additional modeling. Third, summary statistics necessarily compress information. Although our features preserve much of the signal relevant to deep coalescence, they do not retain the full local genealogy or the complete haplotype configuration. This compression is central to the method’s scalability, but it also limits interpretability at individual loci.

A further limitation is that deep population structure and introgression remain intrinsically difficult to separate [35, 63]. Structured ancestral populations can generate old coalescent times and divergent haplotypes, and some recent models have shown that deep structure may explain signals previously interpreted as archaic or ghost admixture. Our updated robustness analyses directly tested this concern by simulating a no-ghost history with deep ancestral structure across the hominin tree. This scenario measurably increased classifier scores, confirming that deep structure is a genuine biological confound. However, it did not produce widespread misclassification: most deep-structure windows were still classified as no-ghost, especially under the strict threshold. Thus, deep ancestral structure should temper interpretation of relaxed-threshold calls, but it does not by itself explain away the high-specificity signal. These alternatives are also not mutually exclusive. Hominin evolution may have involved both long-lived structured populations and episodic admixture among deeply diverged groups. We therefore interpret empirical high-scoring windows as candidate regions requiring downstream validation, with particular attention to local mutation rate, ascertainment, balancing selection, and alternative deep-structure explanations.

Several future directions emerge from this work. First, because DEEP is trained entirely through simulations, it can be readily retrained under alternative demographic models to directly compare competing hypotheses regarding ghost donors, divergent times, and introgression histories. Second, DEEP could be combined with local genealogy inference in a hybrid pipeline using summary-statistic classification as a fast genome-wide screen followed by ARG-based analyses in high-priority candidate regions. Third, additional Denisovan genomes from different times and geographic regions would be essential for resolving weather superarchaic ancestry entered all Denisovan populations or only specific regional populations. More generally, the same general strategy could be extended beyond Denisovan-mediated superarchaic ancestry, including tests for Neanderthal-mediated ghost ancestry, deeper unsampled lineages in other human populations, and analogous questions in nonhuman species where introgression from unsampled or extinct populations is suspected.

In summary, we present DEEP, a scalable framework for detecting candidate regions of Denisovan-mediated superarchaic ancestry without reconstructing genome-wide ARGs. Our results revealed a sparse, externally enriched, and partly shared genomic landscape of superarchaic candidate regions across Oceanians, Tibetans, and Han Chinese. We show that conservative thresholds remain robust to several major no-ghost alternatives, including direct archaic ancestry, sampled-human structure, and substantial deep ancestral structure. At the same time, deep structure, mutation-rate heterogeneity, balancing selection, and ascertainment effects remain important interpretive challenges. More broadly, this work supports the view that human evolutionary history was shaped by interactions among multiple deeply diverged hominin lineages and provides a practical route for studying unsampled ancestry in humans and other species.

## 4 Methods

### 4.1 Analytical coalescent calculations

We derived analytical expectations for the probability that a present-day human window carries Ghost →Denisovan →human ancestry and for the corresponding change in expected human–Denisovan coalescent depth. These calculations were used to evaluate whether the simulation prior included parameter regimes in which ghost ancestry should, in principle, generate a measurable elevation in coalescent depth. We worked in continuous time under the standard coalescent with constant effective sizes within epochs. Calendar times were converted to generations assuming a 25-year generation time.

We evaluated the analytical expressions on a grid spanning Denisovan*→*human migration fractions *m*_D*→*H_ *∈* {0.05, 0.10, 0.15, 0.20, 0.25, 0.30}, ghost*→*Denisovan migration fractions *m*_G*→*D_ ∈ {0.01, 0.02, 0.03, 0.04, 0.05}, human and Denisovan effective sizes *N*_H_, *N*_D_ *∈* {10^3^, 4 *×* 10^3^, 7 *×* 10^3^, 10^4^}, human–Denisovan split times *T*_H|D_ *∈* {300, 500, 700, 900} kya, ghost–Denisovan split times *T*_G|D_ ∈ {1000, 1200, 1400, 1600, 1800} kya, Denisovan*→*human pulse times *T*_D*→*H_ *∈* {40, 60, 80} kya, and ghost*→*Denisovan pulse times *T*_G*→*D_ *∈* {100, 300, 500, 700} kya. We used a present-day human sample size of *n* = 14 haplotypes and an ancestral effective size *N*_A_ = 8000 for deep-time approximations.

Let *A*(*t*) denote the number of ancestral lineages for the *n*-sample at time *t* backwards in time. For *j* ≤*i*≤ *n*, the probability of transitioning from *i* to *j* lineages in time *t*, measured in coalescent units, was computed using the spectral form

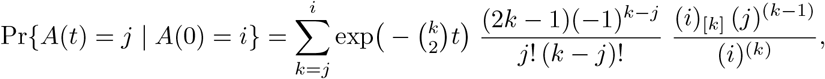

where (*i*)_[*k*]_ = *i*(*i*− 1) … (*i k* + 1) and (*i*)^(*k*)^ = *i*(*i* + 1) … (*i* + *k* −1). We scaled *t* by 2*N*_*e*_ for the corresponding epoch. Conditional on *j* extant lineages at a pulse, the number *K* of lineages receiving migrant ancestry was modeled as

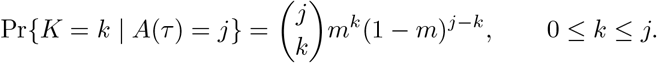

For a Denisovan → human pulse at *T*_D*→*H_, the probability that exactly *m*_*D*_ human ancestral lineages derive from Denisovans was

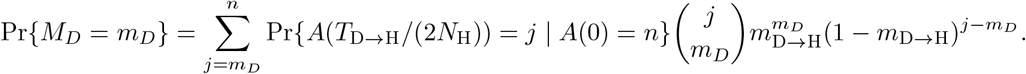

Between the Denisovan*→*human pulse and the ghost*→*Denisovan pulse, Denisovan-derived lineages coalesced at rate 1/(2*N*_D_). Writing Δ = *T*_G*→*D_ *− T*_D*→*H_ in generations, the conditional probability that exactly *m*_*G*_ of the *m*_*D*_ Denisovan-derived lineages trace to the ghost was

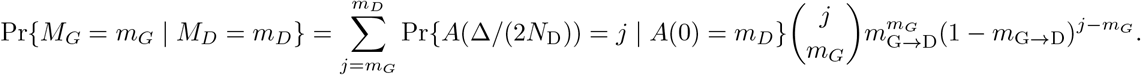

We then summed over *m*_*G*_ *≥* 1 and marginalized over *M*_*D*_ to obtain

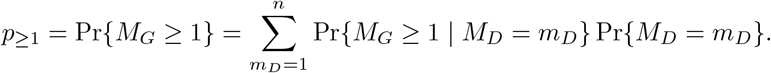

We approximated the expected human–Denisovan coalescent depth with ghost ancestry as a mixture of the human–Denisovan and ghost–Denisovan split depths:

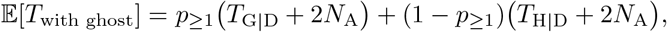

with the no-ghost baseline

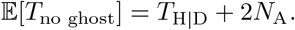

For a simple detectability heuristic, we used Var[*T*_no ghost_] *≈* 2*N*_A_ and flagged parameter settings satisfying

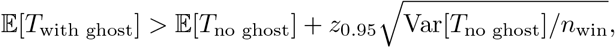

where *z*_0.95_ = 1.644854 and *n*_win_ *∈ {*200, 1400}. For each parameter combination, we recorded Pr*{M*_*D*_ = *m*_*D*_}, Pr*{M*_*G*_ *≥* 1 | *M*_*D*_*}, p*_*≥*1_, *E*[*T*_with ghost_], *E*[*T*_no ghost_], and the detectability indicator.

### 4.2 DEEP neural-network architecture and training

DEEP consists of population-specific neural-network classifiers that assign each retained 50 kb window a probability that its summary-statistic profile was generated under a history in which ancestry from a deeply diverged ghost lineage entered Denisovans and subsequently reached the sampled modern-human population. We trained one classifier for the Papuan application and one classifier for the East Asian application. The East Asian classifier was trained under an East Asian demographic model and was used for downstream analyses of both Tibetan and Han genomes. In both cases, the response variable was defined at the retained-window level: *y* = 1 denoted a window in which at least one retained modern-human haplotype carried Ghost →Denisovan →modern-human ancestry, whereas *y* = 0 denoted a window in which no retained modern-human haplotype carried such ancestry.

For each population-specific classifier, we first merged the feature tables generated from independent simulation batches into a single training table. Candidate input features were restricted to numeric summary-statistic columns whose names began with human or combined. These two prefixes distinguished summaries computed from the phased modern-human haplotypes alone from summaries computed after combining the modern-human and Denisovan samples into diploid genotypes. We did not use all recorded columns as model inputs. Before training, we excluded direct site-count features, singleton-count features, site-frequency-spectrum vector columns, the combined number of individuals, and the combined proportion of segregating sites. This filtering was intended to reduce shortcut learning from mutational opportunity, rare-variant burden, or directly encoded ascertainment differences, while retaining summaries of allele sharing, haplotype divergence, pairwise distances, and Denisovan-conditioned structure.

We standardized features using only the training split. Specifically, we fit a standard scaler to the training feature matrix and then applied the fitted transformation to the validation, test, and empirical feature matrices. We saved both the fitted scaler and the ordered feature list, so that downstream empirical prediction used the same feature ordering and normalization as the simulated training data.

We split the data by simulation bundle rather than by individual row. This was necessary because the simulator generated matched positive and negative examples under the same demographic and mutational draw. A row-wise split would therefore allow closely related examples, or even matched positive–negative counterparts, to appear in different partitions. To avoid this leakage, we assigned each shared simulation bundle to the training, validation, or test partition and then mapped those assignments back to rows. We used an 80/10/10 split into training, validation, and held-out test sets, with stratification by bundle-level class label when possible. The validation set was used for model selection, threshold calibration, and pairwise fine-tuning diagnostics. The test set was kept fully held out until final evaluation.

The classifier was a dense feed-forward neural network with three hidden layers of widths 128, 64, and 32. Each hidden layer used a ReLU activation, *ℓ*_2_ kernel regularization with coefficient 10^−4^, and dropout. Dropout rates were 0.15, 0.10, and 0.05 for the first, second, and third hidden layers, respectively. The output layer consisted of a single sigmoid unit returning

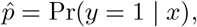

where *x* is the standardized summary-statistic vector for a window.

Training proceeded in two stages. In the first stage, we pretrained the network on all training examples by minimizing binary cross-entropy. We optimized the model with Adam using learning rate 3 ×10^−4^ and batch size 32. During this stage, we monitored validation loss for early stopping, reduced the learning rate when validation loss plateaued, and saved the best checkpoint according to validation ROC–AUC. We allowed up to 1000 epochs, used early stopping with patience 12, reduced the learning rate by a factor of 0.5 after five validation-loss plateau epochs, and imposed a minimum learning rate of 10^−6^. After pretraining, we reloaded the best validation ROC–AUC checkpoint before pairwise fine-tuning.

In the second stage, we fine-tuned the pretrained network on matched positive–negative pairs. For a matched pair, the model produced scores *s*_pos_ and *s*_neg_ for the positive and negative windows, respectively. We defined the paired score difference as

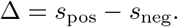

We then optimized a softplus ranking loss with margin 0.02,

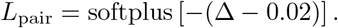

To preserve calibrated binary classification while encouraging the positive member of each matched pair to receive a higher score, we combined the pair-ranking loss with a binary cross-entropy term and the usual regularization penalty:

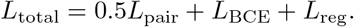

We optimized this objective with Adam using learning rate 10^−4^, batch size 128, and a maximum of 150 epochs. After each epoch, we evaluated matched-pair performance on the validation set. We retained the weights that maximized validation pairwise accuracy, breaking ties by the mean positive-minus-negative score difference. Training stopped when validation pairwise performance failed to improve for 15 epochs.

After training, we defined operating thresholds using the negative-score distribution in the set-aside test split. During model development, we monitored the behavior of low-FPR operating points in the validation set, but the empirical analyses used the corresponding thresholds as defined in the held-out test data. Because genome-wide empirical scans involve tens of thousands of 50 kb windows and the target signal is expected to be rare, we emphasized false-positive-rate control rather than a single accuracy-maximizing threshold.

We used two test-set FPR thresholds. The more conservative threshold targeted an approximately genome-wide false-positive rate of 1/55,000, corresponding to roughly one expected false positive across an autosomal genome-wide scan. This threshold defined the highest-confidence set of candidate windows. We also used a less conservative threshold targeting a false-positive rate of 1/1,000. This broader threshold increased power to detect weaker or more fragmented Ghost→ Denisovan →modern-human signals while still imposing stringent false-positive control. We treated these thresholds as complementary: the 1/55,000 threshold prioritized specificity, whereas the 1/1,000 threshold provided a larger discovery set for analyses of regional clustering, cross-population sharing, and overlap with external annotations.

The test split was not used for gradient updates, early stopping, or model selection. It was reserved until after training to estimate final model performance and to define the empirical operating thresholds. We retained threshold-specific validation and test summaries, row-level predictions, split assignments, fitted scalers, ordered feature lists, pairwise diagnostics, and summary metrics for downstream analyses.

### 4.3 Simulated Data Sets Used for Training and Testing

We generated supervised training data with `msprime` [64–67] by simulating 50 kb autosomal windows under population-specific demographic models that included modern humans, Neanderthals, Denisovans, and a deeply diverged unsampled ghost lineage. We trained one model for Papuan application and one structurally parallel model for East Asian application. The East Asian model was applied to both Tibetan and Han genomes. For each population-specific classifier, all training examples were drawn from a single demographic model. Positive and negative simulations shared the same demographic structure, mutationrate model, sampling scheme, and filtering procedure; they differed only in whether the Ghost → Denisovan pulse was present.

We parameterized time in generations and used a 25-year generation time when converting calendar dates to model time. Each ancestry simulation used a sequence length of 50 kb, a recombination rate of 1 × 10^−8^ per base per generation, and migration recording enabled. After ancestry simulation, we overlaid neutral binary mutations using a window-specific mutation rate. Each retained window ultimately contained 16 haplotypes: 14 present-day modern-human haplotypes from the target population and two haplotypes from one sampled Denisovan diploid. The Denisovan diploid was sampled at approximately 63.9 kya, corresponding to the midpoint of the estimated age interval for the high-coverage Denisovan genome.

The Papuan and East Asian models shared the same deeper hominin scaffold. Described backward in time, the sampled non-African population first merged into the out-of-Africa lineage. The out-of-Africa lineage and the African lineage then merged into an ancestral modern-human lineage at 140 kya. This ancestral modern-human lineage entered a deeper modern-human ancestral population at 220 kya. The Denisovan and Neanderthal lineages merged within the archaic branch at a time drawn from

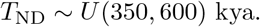

The modern-human ancestral branch and the archaic branch then merged at 600 kya. Finally, the ghost lineage was constrained to split more deeply than this modern-human–archaic divergence. Thus, the two population-specific models differed in the terminal non-African branch and in target-population-specific parameter ranges, but not in the basic relationship among modern humans, Neanderthals, Denisovans, and the ghost lineage.

The population-specific part of the model occurred on the terminal non-African branch. In the Papuan model, the sampled modern population split from the out-of-Africa lineage at

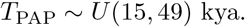

In the East Asian model, the sampled modern population split from the out-of-Africa lineage at

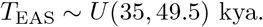

In both models, the out-of-Africa split time was drawn from

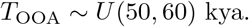

Both population-specific models included a bottleneck on the sampled modern-human lineage. This bottleneck was included in both positive and negative simulations and therefore formed part of the shared demographic background rather than the class-defining signal.

We drew effective population sizes from broad ranges intended to capture uncertainty in the relevant demographic histories. In both models, the Denisovan and Neanderthal effective population sizes were drawn from *U* (1000, 4500), the ghost effective population size from *U* (1000, 10000), and the archaic ancestral effective population size from *U* (1000, 6000). The African and out-of-Africa ancestral sizes were fixed at 12,300 and 2,100, respectively. The sampled modern-human population size was drawn separately for each population-specific model. In the Papuan model, the present-day target-population size was drawn from *U* (2500, 3700). In the East Asian model, it was drawn from *U* (6000, 10000). These population-size parameters were sampled independently across simulation bundles.

All simulations included Denisovan-to-modern-human and Neanderthal-to-modern-human introgression. In the Papuan model, the Denisovan admixture fraction was drawn from

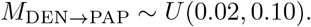

In the East Asian model, the Denisovan admixture fraction was drawn from

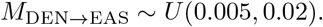

In both models, the Neanderthal admixture fraction was drawn from

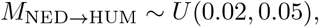

where HUM denotes the sampled modern-human population in the corresponding model. The Denisovan-to-modern-human and Neanderthal-to-modern-human pulse times were drawn uniformly between the split time of the sampled modern population and the out-of-Africa split. Although these events were implemented in msprime as backward-time mass migrations, we describe them here in their corresponding forward-time direction.

Positive and negative examples differed only in whether ghost ancestry entered the Denisovan lineage. For positive windows, the Ghost*→*Denisovan admixture fraction was drawn from

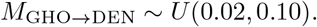

For negative windows, this fraction was set to zero. The Ghost→Denisovan pulse time was drawn uniformly between the sampled Denisovan age and the Neanderthal–Denisovan split. This construction produced the target signal of interest: ancestry from an unsampled deeply divergent lineage entering Denisovans and subsequently reaching modern humans through Denisovan introgression.

For each simulated window, we first sampled an oversized modern-human panel together with the Denisovan diploid. The oversized panel contained 20 modern-human diploids from the target population. We then removed singleton sites before final downsampling. Singleton status was defined across all oversampled modern-human haplotypes, not across the final retained sample. Specifically, we removed sites with minor allele count one among the oversampled modern-human haplotypes. Only after this singleton-filtering step did we downsample whole diploid pairs to the final panel of seven modern-human diploids. This ordering ensured that rare-site filtering reflected the larger simulated sample rather than stochastic rarity induced by the final downsampling step.

We accepted positive examples only when the final retained panel contained Ghost → Denisovan→ modern-human ancestry. Specifically, at least one of the final 14 modern-human haplotypes had to carry ghost-derived ancestry that had passed through the Denisovan lineage. We accepted negative examples only when none of the final 14 modern-human haplotypes carried such ancestry. These criteria ensured that the label referred to the retained analysis panel rather than to ancestry present only in discarded oversampled individuals.

We assigned labels from recorded ancestry rather than from simulation parameters alone. For each tree sequence, we scanned the recorded migration table for genomic segments in which a Denisovan lineage moved into the ghost population backward in time, corresponding to forward-time Ghost*→*Denisovan ancestry. For each such segment, we evaluated the local tree at the midpoint of the segment and identified descendant sampled nodes. We then counted the number of retained modern-human haplotypes descended from those migration segments. A retained window was labeled positive if this count was at least one and negative if this count was zero. We also recorded the number of ghost-ancestry tracts reaching sampled humans, the number of retained human haplotypes carrying ghost ancestry, and whether the sampled Denisovan carried ghost ancestry.

To reproduce empirical variation in the number of retained sites across 50 kb windows, we fit a population-specific mutation-rate heterogeneity model before generating the final supervised training data. We fit this model separately for the Papuan and East Asian training pipelines, using the same sampling scheme, singleton-filtering order, and retained-site definition used by the corresponding simulator. The goal of this fitting step was not to estimate a genome-wide biological mutation rate in isolation, but to make simulated windows match the empirical distribution of retained sites after the same filtering and feature-extraction steps used downstream.

For each population-specific fit, we used empirical autosomal windows processed through the corresponding feature-extraction pipeline and fit to the column combined num sites. This count records the number of sites retained in a 50 kb window after allele re-encoding and restriction to sites polymorphic among the 14 retained modern-human haplotypes. We included windows with zero retained sites during fitting, so that the fitted model captured both low-diversity and high-diversity regions of the empirical distribution.

We modeled the mutation rate in window *w* as

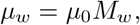

where *µ*_0_ is a baseline mutation-rate parameter and *M*_*w*_ is a window-specific multiplicative rate modifier. We modeled *M*_*w*_ as a gamma-distributed random variable with mean one,

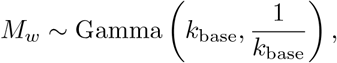

where *k*_base_ controls the degree of among-window mutation-rate heterogeneity. Under this parameterization,

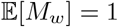

and

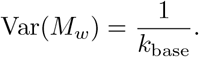

Thus, *µ*_0_ controls the average mutation rate across windows, whereas *k*_base_ controls the dispersion of mutation rates among windows. Smaller values of *k*_base_ allow greater heterogeneity in retained site counts, while larger values concentrate window-specific rates more tightly around *µ*_0_.

We fit the parameters by direct simulation and grid search. For each candidate pair (*µ*_0_, *k*_base_), we drew a mutation rate for each replicate window, simulated no-ghost windows with msprime under the corresponding population-specific model, and applied the same high-level processing order used in the training simulator. This order was: simulate an oversampled modern-human panel together with one Denisovan diploid; overlay mutations; remove singleton sites defined across all oversampled modern-human haplotypes; downsample whole diploid pairs to the final seven modern-human diploids; re-encode alleles by minor/major state; and count sites that remained polymorphic among the retained modern-human haplotypes. Because all candidate simulations used no-ghost histories, the fitted mutation model matched the empirical retained-site-count distribution without introducing the class-defining Ghost*→*Denisovan signal.

For the Papuan mutation model, we fit to the empirical Papuan autosomal feature table. Candidate simulations used 50 kb windows, recombination rate 1 × 10^−8^, a 25-year generation time, seven retained Papuan diploids, one Denisovan diploid, and minor-allele singleton filtering across the oversampled Papuan haplotypes. During this fitting step, the oversampled Papuan panel contained 50 diploids before downsampling. We evaluated

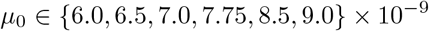

and

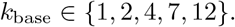

For each grid point, we simulated 400 no-ghost replicate windows. The best-fitting Papuan model used

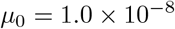

and

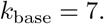

Thus, the Papuan training simulations drew window-specific mutation-rate multipliers from

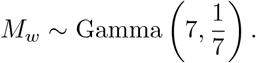

For the East Asian mutation model, we fit to the empirical East Asian autosomal feature table using the same retained-site definition. Candidate simulations used 50 kb windows, recombination rate 1 × 10^−8^, a 25-year generation time, seven retained East Asian diploids, one Denisovan diploid, and minor-allele singleton filtering across the oversampled East Asian haplotypes. During this fitting step, the oversampled East Asian panel contained 20 diploids before downsampling. We evaluated

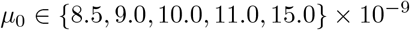

and

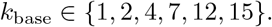

For each grid point, we simulated 800 no-ghost replicate windows. The best-fitting East Asian model used

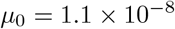

and

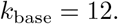

Thus, the East Asian training simulations drew window-specific mutation-rate multipliers from

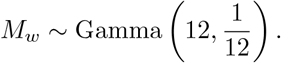

We selected the best-fitting parameter combination by comparing the simulated and empirical retained-site-count distributions. Let *Q*_sim,*θ*_(*p*) and *Q*_emp_(*p*) denote the simulated and empirical quantiles at probability *p* for candidate parameter vector *θ*. The primary loss was a relative squared quantile-grid loss,

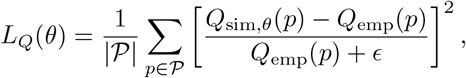

where *ϵ* is a small numerical constant. We used a dense quantile grid spanning the empirical distribution from the 0.01 through 0.99 quantiles, together with the 0.999 quantile. We set the tail-weighting parameter to zero, so the quantile grid emphasized the overall retained-site-count distribution rather than disproportionately weighting the extreme upper tail.

For the Papuan fit, the objective placed all weight on the quantile-grid loss. For the East Asian fit, the objective combined the quantile-grid loss with additional relative squared penalties on the mean and median retained-site counts. These additional terms stabilized the East Asian fit while retaining the quantile grid as the main constraint on the full distributional shape.

The fitted models closely matched the central and upper-central portions of the empirical retained-site-count distributions, which were the portions most relevant for generating realistic training examples. In the Papuan fit, the empirical median, 75th percentile, 90th percentile, and 99th percentile of retained site counts were 28, 42, 58, and 97, respectively; the corresponding simulated values under the selected model were 24, 41, 55, and 96. In the East Asian fit, the empirical median, 75th percentile, 90th percentile, and 99th percentile were 37, 53, 71, and 118, respectively; the corresponding simulated values were 36, 53.25, 71, and 108. The most extreme empirical tail contained a small number of high-count windows that were not the primary target of the fit, because the final objective did not upweight the extreme tail.

The selected Papuan and East Asian mutation-rate models were written to population-specific configuration files and were then used by the corresponding training simulators to draw one mutation rate per simulated window.

Whenever possible, the simulator generated matched positive and negative examples under the same drawn demographic and mutational parameters. In each matched pair, the positive and negative windows shared the same nuisance draw and differed only in whether the Ghost → Denisovan pulse was active. This paired design helped separate the signal of ghost-mediated Denisovan ancestry from background variation in mutation rate, archaic admixture proportions, split times, and population sizes. We propagated the bundle identifiers and pair labels into the downstream feature table so that matched examples could be kept within the same train, validation, or test partition and used during pairwise neural-network fine-tuning.

For each population-specific training run, we generated independent simulation batches on a compute cluster. Each batch targeted 50,000 accepted positive windows and 50,000 accepted negative windows before downstream feature extraction and filtering of windows lacking retained human-polymorphic sites. The per-batch genotype matrices and metadata were written separately for positive and negative examples and were later converted into summary-statistic feature tables for model training.

### 4.4 Feature Extraction

We converted each retained simulated 50 kb window into a vector of summary statistics. Each genotype matrix contained physical positions in the first column, followed by 14 present-day modern-human haplotypes and two Denisovan haplotypes. Before computing features, we re-encoded each segregating site so that the minor allele in the retained 16-haplotype panel was coded as 1 and the major allele was coded as 0. We then restricted analysis to sites that remained polymorphic among the 14 modern-human haplotypes. The same human-polymorphic site mask was used for both the human-only and combined feature blocks. Windows with no retained human-polymorphic sites were discarded.

For every retained window, we computed two complementary sets of summary statistics. The first set, denoted with the human prefix, summarized the 14 modern-human haplotypes as phased haploid sequences. The second set, denoted with the combined prefix, summarized an eight-individual diploid panel formed by collapsing the 14 modern-human haplotypes into seven modern-human diploids and the two Denisovan haplotypes into one Denisovan diploid.

For the human-only block, we computed the number of retained sites and haplotypes, the mean and variance of the sitewise allele-frequency distribution, mean heterozygosity, the proportion of segregating sites, the site-frequency spectrum, Tajima’s *D*, and the number of singletons among retained human-polymorphic sites. We also summarized pairwise haplotype divergence by computing all pairwise Hamming distances among the 14 modern-human haplotypes and recording their mean, variance, skewness, kurtosis, and the five largest observed pairwise distances. These summaries were designed to capture shifts in local coalescent depth and heterogeneity in haplotype divergence within each window.

We additionally computed Denisovan-conditioned haplotype features on the same retained human-polymorphic sites. These features quantified how modern-human variation aligned with the sampled Denisovan haplotypes. First, we identified sites at which at least one Denisovan haplotype carried the minor allele and at least one modern-human haplotype also carried that allele. For these Denisovan-shared sites, we recorded the total number of shared sites and stratified them by the number of modern-human haplotypes carrying the shared allele: one haplotype, two haplotypes, three to four haplotypes, five to seven haplotypes, or at least eight haplotypes. We also recorded the mean and variance of the modern-human allele count conditional on Denisovan sharing.

Second, we measured the affinity between each modern-human haplotype and the sampled Denisovan haplotypes. For each modern-human haplotype, we computed its Hamming distance to each of the two Denisovan haplotypes across retained sites and kept the smaller of the two distances. We summarized the resulting 14 human-to-Denisovan distances by their minimum, mean, variance, lower-tail quantile, and the three closest haplotype-to-Denisovan distances.

Third, we summarized how Denisovan-shared alleles were distributed across the modern-human haplotypes. For each modern-human haplotype, we counted the number of Denisovan-shared alleles it carried. We then recorded the mean, variance, skewness, kurtosis, maximum, Gini coefficient, and entropy of this per-haplotype distribution. These features were intended to distinguish diffuse allele sharing from sharing concentrated on a small number of haplotypes, as expected for tract-like introgressed ancestry.

Fourth, we summarized the spatial clustering of Denisovan-shared sites within each 50 kb window. We computed nearest-neighbor distances among Denisovan-shared sites and recorded their mean, variance, skewness, and kurtosis. We also divided the window into 5 kb bins and recorded the maximum number of Denisovan-shared sites in any bin and the number of occupied bins. Finally, we computed run-like summaries of Denisovan-shared alleles on each modern-human haplotype, including the longest run measured in sites and base pairs, the number of runs per haplotype, and the mean run size per haplotype. We summarized these run statistics across haplotypes by their means, variances, and maxima.

For the combined block, we collapsed the phased modern-human haplotypes into seven diploid genotypes and collapsed the two Denisovan haplotypes into one Denisovan diploid genotype. Using this eight-individual diploid matrix, we computed the number of retained sites and individuals, the mean and variance of the sitewise allele-frequency distribution, expected and observed heterozygosity, the proportion of segregating sites, the site-frequency spectrum, Tajima’s *D*, and the number of singletons. We also computed all pairwise city-block distances among the eight diploid genotypes and recorded their mean, variance, skewness, kurtosis, and the five largest values.

To capture within-window heterogeneity, we divided each 50 kb window into 10 kb subwindows. For each diploid individual in the combined panel, we computed the proportion of heterozygous sites in each occupied subwindow. We then calculated all pairwise absolute differences among occupied subwindows within each individual and summarized the resulting distribution by its mean, variance, skewness, kurtosis, and five largest values. These summaries were included to capture local clustering of variation within windows, which may arise when introgressed ancestry is concentrated over part of a window rather than distributed uniformly.

The feature-extraction step also carried forward simulation metadata, including the binary label, bundle identifier, pair role, demographic parameters, mutational parameters, and ancestry-derived quantities used to define the label. These metadata columns were not treated as neural-network inputs. They were retained to support grouped data splitting, matched-pair fine-tuning, diagnostic analyses, and reproducibility of the simulation pipeline.

### 4.5 DEEP identifies candidate ghost-introgressed windows

We used the genealogical signal described above to train DEEP classifiers for window-level detection of Ghost →Denisovan →human ancestry. For each simulated 50-kb window, the underlying tree sequence records ancestry of every haplotype, therefore we label windows accordingly whether it contains ghost ancestry transmitted through Denisovans into humans. The neural network was trained only on observable window-level summary statistics computed from the simulated genotype data.

We trained separate classifiers for the Oceanian and East Asian analyses so that demographic background, sampling scheme, and nuisance variation matched the empirical populations to which each classifier would later be applied. In both cases, simulations with known ancestry labels were used to generate training data, followed by feature extraction, scaling, supervised model training, and for prediction on empirical genomic windows (Fig. 4a). The resulting score for each window can be interpreted as a model-based measure of support for ghost-mediated ancestry under the corresponding simulation model.

This framework separates the definition of the target state from the inference: simulations define whether a window contains Ghost →Denisovan →human ancestry, whereas the classifier learns which observable patterns of human variation, human–Denisovan sharing, and local haplotype relationships are informative about that state. Empirical windows receiving high scores are therefore interpreted as candidates whose summary-statistic profiles resemble simulated ghost-positive windows.

The simulation-trained workflow was computationally tractable at genome scale. Genotype simulation and summary-statistic construction were parallelized as 25-task arrays for each population-specific model, followed by one final neural-network training. Across all population-specific models, the complete training required 436.0 CPU-hours (Table 1), with most computation effort devoted to generating labeled simulated genotype data. Because empirical application uses window-level summary statistics rather than genome-wide ARG reconstruction, DEEP provides a scalable framework for genome-wide screening of superarchaic ancestry.

### 4.6 Processing empirical modern-human and Denisovan genomes

We applied the trained classifiers to empirical genotype data from three modern-human target populations: Oceanians, Tibetans, and Han Chinese. The Oceanian data were drawn from the Simons Genome Diversity Project (SGDP) [46], the Tibetan data from Lu et al.[47], and Han Chinese data from the high-coverage 1000 Genomes Project/IGSR genomes [48–50]. All empirical processing was performed on autosomes 1– 22 in GRCh37/hg19 coordinates. For each population, we merged the modern-human genotypes with a filtered high-coverage Denisovan VCF [2], removed singleton sites using a larger population-specific panel, extracted the final high-coverage samples used for prediction, applied a hard genomic mask and minor-allele-frequency filter, and converted the resulting VCFs into the same 50 kb window-level summary-statistic representation used for simulated training data.

The empirical singleton-filtering procedure paralleled the simulation pipeline. In the simulations, singleton removal was performed in an oversampled modern-human panel before downsampling to the final retained samples. We therefore defined empirical singletons in a larger population-specific panel rather than in the final prediction panel alone. The singleton-definition panels contained 19 Oceanian diploids, corresponding to all Oceanians available in SGDP [46]; 38 Tibetan diploids from Lu et al. [47]; and 108 CHB diploids from the high-coverage 1000 Genomes/IGSR data set. This larger-panel filtering reduced the chance that variants would be treated as rare only because of undersampling in the smaller finalset of high-coverage individuals used for prediction.

For each chromosome, we first prepared a modern-human input VCF or BCF and the corresponding filtered Denisovan VCF. The Oceanian pipeline used SGDP BCFs, which were rewritten as bgzipped VCFs before merging. The Tibetan and Han pipelines used phased hg19 VCFs. For the Tibetan and Han inputs, we checked whether the VCF header contained an INFO/END definition and added this definition when necessary before reheadering, recompressing, and indexing the file. We then indexed the Denisovan VCF and merged the full modern-human chromosome file with the Denisovan VCF using bcftools merge with multiallelic merging disabled and missing genotypes set to reference.

After merging, we identified which samples from the population-specific singleton-definition panel were present in the merged VCF. For Oceanians, we intersected the full Oceanian sample list with the merged VCF sample list. For Tibetans, we generated the complete sample list and intersected it with the merged VCF sample list. For Han, we intersected the CHB sample list with the merged VCF sample list. We also extracted the Denisovan sample column or columns directly from the Denisovan VCF. The singleton-definition panel for each chromosome consisted of the available population-specific modern-human samples together with the Denisovan sample column or columns.

We constructed the final prediction panel separately. For each population, the final sample list consisted of the selected high-coverage modern-human samples plus the Denisovan sample. The pipeline required all final samples to be present in the merged VCF and stopped if any requested final sample was missing. This prevented chromosome-specific sample dropping and ensured that all empirical windows were represented with a consistent sample layout for downstream feature extraction and prediction.

Singleton filtering was performed before final sample extraction. We subset the merged VCF to the singleton-definition panel, retained only biallelic SNPs, and recomputed allele count and allele number tags in that panel. We then retained only sites whose minor allele count was at least two in the singleton-definition panel. Operationally, this filter retained sites satisfying

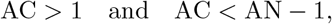

where AC and AN were recomputed after subsetting to the singleton-definition panel. This removed alternate-allele singletons, reference-allele singletons at nearly fixed alternate sites, and monomorphic records introduced by subsetting. We then extracted the final high-coverage modern-human samples plus Denisovan from the full merged VCF and intersected this final-sample VCF with the non-singleton site set defined in the larger panel. The intersection was written from the final-sample VCF, preserving the final sample columns while enforcing the larger-panel singleton filter. We then recomputed AC and AN in the final sample set and indexed the resulting VCF.

We next applied a hard genomic mask in hg19 coordinates [68–72]. The mask excluded low-mappability regions, assembly gaps, CpG islands, RepeatMasker-annotated repeats, all CpG dinucleotides, and ENCODE blacklist regions. Low-mappability regions were defined from the CRG 100-mer mappability track by masking intervals with mappability less than 1.0. Assembly gaps were obtained from the UCSC gap table. CpG islands were obtained from the UCSC `cpgIslandExt` table. Repeat annotations were obtained from the UCSC RepeatMasker rmsk table. We additionally scanned the hg19 reference FASTA directly and masked every CpG dinucleotide as a two-base-pair interval. All mask components were sorted, merged, compressed, and indexed.

For each chromosome, we excluded all variants overlapping the combined hard mask. We then recomputed minor allele frequencies with`bcftools +fill-tags` and retained variants satisfying

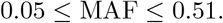

The lower bound removed rare variants outside the feature-distribution regime targeted by the classifier. The upper bound was set slightly above 0.5 so that all common biallelic variants could be retained after tag recomputation. The resulting masked and MAF-filtered VCFs were compressed, indexed, and used as the input to empirical window construction.

We converted each filtered chromosome-level VCF into haplotype matrices for non-overlapping 50 kb windows. For each VCF record, we wrote the physical position in the first column. We then split each phased modern-human diploid genotype into two allele columns. The final empirical prediction panel therefore contained 14 modern-human haplotype columns, corresponding to seven modern-human diploids, followed by two Denisovan allele columns from one Denisovan diploid. Because the Denisovan genotype was treated as an unphased diploid genotype, the order of the two Denisovan allele columns was arbitrary and was used only to represent the two allelic states at each site. Monomorphic all-reference rows were skipped. For each chromosome, non-empty window matrices were saved in a compressed `npz` file, with each matrix keyed by its window coordinates.

We then computed empirical summary statistics from these window-level haplotype matrices using the same feature-extraction functions used for simulated data. Within each window, we re-encoded sites so that the minor allele in the retained human–Denisovan panel was coded as 1 and the major allele as 0. We then restricted the analysis to sites polymorphic among the retained modern-human haplotypes. Windows with no retained human-polymorphic sites were omitted from the prediction table. For each remaining window, we computed the human-only phased-haplotype feature block and the combined diploid feature block, including the Denisovan-conditioned haplotype features described above. We wrote one feature table per chromosome and then concatenated chromosomes 1–22 into a single autosomal empirical feature table for each target population.

Finally, we scored empirical windows with the population-matched classifier. The Papuan-trained classifier was applied to the Oceanian data. The East Asian-trained classifier was applied to both Tibetan and Han data. Before prediction, empirical feature matrices were restricted to the ordered feature list saved during training and standardized with the scaler fit on the corresponding simulated training split. The trained neural network then produced a predicted probability for each empirical 50 kb window, which was compared to the operating thresholds defined from the held-out simulated test split.

## 5 Acknowledgments

XZ was supported by the National Institutes of Health/National Institute of General Medical Sciences under grant R35GM154856. SZ was supported by the National Institutes of Health grant R01HG005855. NM was supported by the National Human Genome Research Institute under grant T32-HG000040.

## 6 Author Contributions

XZ conceived the study. SZ and NM developed the analytical framework. NM performed all analyses and implemented the machine learning framework. NM wrote the manuscript. XZ and SZ contributed to manuscript revision. All authors approved the final manuscript.

## 7 Data Availability

The data used in this study are available from publicly accessible sources as described in the manuscript and supplementary materials. Processed data and intermediate files generated during the current study are available from the corresponding authors upon reasonable request.

## 8 Code Availability

Code used to perform the simulations, train the classifiers, generate empirical predictions, and reproduce the analyses is available from the corresponding authors upon reasonable request.

## 10 Appendix

### 10.1 Additional Summary Statistic Distributions

### 10.2 Robustness profile definitions

All robustness profiles were simulated as no-ghost negative controls. In every profile, the Ghost → Denisovan pulse was disabled by setting the ghost-to-Denisovan introgression proportion to zero. The profiles therefore differ from the corresponding population-specific no-ghost base model only through nuisance parameters, through added sampled-human structure, or through added deep ancestral structure across the hominin tree. Uniform draws are denoted *U* (*a, b*). Direct archaic ancestry values are pulse proportions in the msprime mass-migration model.

Simple profiles retained the population graph and sampling scheme of the corresponding population-specific simulator. The Oceanian profiles perturbed the Papuan no-ghost model, whereas the East Asian profiles perturbed the East Asian no-ghost model. Structured sampled-human profiles added two sampled human subpopulations, denoted A and B, before singleton filtering and final downsampling. The deep ancestral-structure profile retained a single sampled human population but replaced the immediate deep hominin ancestor with two ancestral demes, one on the human side of the tree and one on the archaic side of the tree, with low migration before their common ancestry.

#### 10.2.1 Simple Oceanian no-ghost profiles

In the Oceanian simple profiles, direct Denisovan ancestry refers to the Denisovan-to-Papuan pulse proportion, and direct Neanderthal ancestry refers to the Neanderthal-to-Papuan pulse proportion. The fitted mutation model refers to the population-specific window-level mutation-rate mixture used in the main simulation workflow.

When the Oceanian recent bottleneck mode was active, the bottleneck size was drawn from *U* (20, 80), and the intermediate post-bottleneck size was drawn from *U* (150, 600). Human-polymorphism-enriched site retention used five site-class-specific retention probabilities: human-polymorphic sites not shared with Denisovan were retained with probability *U* (0.90, 1.00); human-polymorphic sites also present in Denisovan were retained with probability *U* (0.80, 0.95); Denisovan-only sites were retained with probability *U* (0.10, 0.30); sites fixed alternate in humans were retained with probability *U* (0.40, 0.70); and all other sites were retained with probability *U* (0.35, 0.60).

#### 10.2.2 Simple East Asian no-ghost profiles

In the East Asian simple profiles, direct Denisovan ancestry refers to the Denisovan-to-East-Asian pulse proportion, and direct Neanderthal ancestry refers to the Neanderthal-to-East-Asian pulse proportion.

When the East Asian recent bottleneck mode was active, the bottleneck size was set to 100, and the intermediate post-bottleneck size was set to one half of the drawn East Asian effective population size. Human-polymorphism-enriched site retention used the same five retention classes as in the Oceanian profiles: *U* (0.90, 1.00) for human-polymorphic sites, *U* (0.80, 0.95) for human-polymorphic sites also present in Denisovan, *U* (0.10, 0.30) for Denisovan-only sites, *U* (0.40, 0.70) for sites fixed alternate in humans, and *U* (0.35, 0.60) for all other sites.

#### 10.2.3 Structured sampled-human no-ghost profiles

Structured sampled-human profiles replaced the single sampled human population with two sampled sub-populations, A and B. For the Oceanian model, A and B merged into a shared Papuan parent; for the East Asian model, A and B merged into a shared East Asian parent. The structured parent then followed the corresponding base model into the deeper population graph. Direct Denisovan and Neanderthal introgression were applied separately to A and B. All structured sampled-human profiles set the ghost-to-Denisovan introgression proportion to zero.

The tables below list only the parameters introduced by the structured sampled-human profiles. Final A:B gives the number of final diploid individuals sampled from subpopulations A and B. Oversample A:B gives the diploid mixture simulated before singleton filtering and final downsampling. The substructure split time is reported in thousands of years before present.

For the structured bottlenecked-minority profile, the bottleneck was applied only to subpopulation B. The bottleneck size was drawn from *U* (35, 150). For the structured callability-bias profile, human-polymorphic sites were retained with probability *U* (0.90, 1.00); human-polymorphic sites also present in Denisovan were retained with probability *U* (0.80, 0.98); Denisovan-only sites were retained with probability *U* (0.10, 0.30); sites fixed alternate in humans were retained with probability *U* (0.35, 0.70); and all other sites were retained with probability *U* (0.30, 0.60).

#### 10.2.4 Deep ancestral-structure no-ghost profile

The deep ancestral-structure profile was designed to test a more direct biological alternative to Ghost → Denisovan introgression. Unlike the structured sampled-human profiles above, this profile did not divide the present-day sampled humans into A and B subpopulations. Instead, it retained a single sampled human population and introduced substantial structure deeper in the hominin tree. After the AMH-side and archaic-side lineages were formed, they entered two distinct ancestral demes, a deep human-side ancestor and a deep archaic-side ancestor, before merging into a common deep ancestral root. Low symmetric migration was allowed between the two deep ancestral demes during the structured phase. This creates no-ghost windows in which human–Denisovan coalescence can be deep because of ancestral structure rather than because of superarchaic introgression into Denisovans. Schematic demesdraw representations of the Oceanian and East Asian versions of this no-ghost model are shown in Supplementary Figure 7.

For the Oceanian version, the sampled human population was the Papuan parent population used by the Oceanian simulator. For the East Asian version, the sampled human population was the East Asian parent population used by the East Asian simulator. Modern humans were still allowed to receive direct Denisovan and Neanderthal ancestry, but the Ghost*→*Denisovan pulse was set to zero in all replicates.

This profile was included in the main robustness set because it targets the principal genealogical confound for the classifier: long human–Denisovan coalescence times generated by ancestral population structure rather than by a Ghost*→*Denisovan*→*human path.

#### 10.2.5 Shared processing of robustness profiles

All robustness profiles were simulated as 50-kb windows with recombination rate 1*×* 10^−8^ and binary mutations. Each accepted replicate contained seven final modern human diploids and one Denisovan diploid. For simple profiles, singleton filtering was applied to the oversampled human panel before down-sampling to seven diploids. For structured sampled-human profiles, singleton filtering was applied across all oversampled A and B human haplotypes before choosing the final A:B diploid mixture. For the deep ancestral-structure profile, singleton filtering was applied to the oversampled single human parent population before downsampling to seven diploids. In all cases, singleton filtering used the oversample-defined minor-allele criterion.

The resulting genotype matrices were converted to the same human-only and combined summary-statistic feature sets used by the trained classifiers. The saved feature list and saved scaler for the corresponding model family were then applied before prediction. Because all robustness profiles set the ghost-to-Denisovan introgression proportion to zero, the proportion of windows exceeding a classifier threshold estimates the no-ghost false-positive call rate under that robustness profile.

### 10.3 Scenario-level robustness results

### 10.4 Tibetan and Han Overlap with Private Variant Tracts

